# How to train your neuron: Developing a detailed, up-to-date, multipurpose model of hippocampal CA1 pyramidal cells

**DOI:** 10.64898/2026.03.19.712861

**Authors:** Luca Tar, Sára Sáray, Máté Mohácsi, Tamás F. Freund, Szabolcs Káli

## Abstract

Anatomically and biophysically detailed models of neurons have been widely used to study information processing in these cells. Most studies focused on understanding specific phenomena, while more general models that aim to capture various cellular processes simultaneously remain rare even though such models are required to predict neuronal behavior under more complex, natural conditions.

In this study, we aimed to develop a detailed, data-driven, general-purpose biophysical model of hippocampal CA1 pyramidal neurons. We leveraged extensive morphological, biophysical and physiological data available for this cell type, and established a systematic workflow for model construction and validation that relies on our recently developed software tools. The model is based on a high-quality morphological reconstruction and includes a diverse curated set of ion channel models. After incorporating the available constraints on the distribution of ion channels, the remaining free parameters were optimized using the Neuroptimus tool to fit a variety of electrophysiological features extracted from somatic whole-cell recordings. Validation using HippoUnit confirmed the model’s ability to replicate key electrophysiological features, including somatic voltage responses to current input, the attenuation of synaptic potentials and backpropagating action potentials, and nonlinear synaptic integration in oblique dendrites.

Our model also included active dendritic spines, modeled either explicitly or by merging their biophysical mechanisms into those of the parent dendrite. We found that many aspects of neuronal behavior were unaffected by the level of detail in modeling spines, but modeling nonlinear synaptic integration accurately required the explicit modeling of spines.

Our data-driven model of CA1 pyramidal cells matching diverse experimental constraints is a general tool for the investigation of the activity and plasticity of these cells and can also be a reliable component of detailed models of the hippocampal network. Our systematic approach to building and validating general-purpose models should apply to other cell types as well.

**Author Summary:** The brain processes information through the activity of billions of individual neurons. To understand how these cells work, scientists build detailed computer models that reproduce their electrical behavior. These models make it possible to explore situations that are difficult or impossible to test experimentally. However, many existing neuron models were designed to explain only a few specific phenomena, which limits their usefulness in more complex settings.

In this study, we developed a comprehensive computer model of a hippocampal CA1 pyramidal neuron, a cell type that plays a central role in learning and memory. We built the model using extensive experimental data and applied automated methods to ensure that it reproduces a broad range of observed neuronal behaviors.

We also examined how small structures called dendritic spines—tiny protrusions where most synaptic communication occurs—affect how neurons combine incoming signals. We found that even simplified models without individual spines can capture many aspects of neuronal activity, but understanding more complex forms of signal integration requires modeling spines explicitly. Our work also supports the development of more realistic simulations of brain circuits.

## Introduction

In the last few decades, the need for anatomically and biophysically detailed neuronal models in neuroscientific research has increased massively. Such models can be used to examine the function and behavior of specific types of neurons, to explore the various mechanisms behind the observed electrophysiological behavior, and to predict neuronal behavior and function under circumstances that are difficult to study directly in experiments [1] [2] [3] [4] [5] [6] [7] [8]. In addition, such detailed cellular models are the fundamental building blocks of detailed network models [9] [10] [11] [12] [9] [13].

Recently, it has become standard to support new experimental findings with models that are tuned to mimic the observed behavior, but many of the models in earlier studies had important limitations. In particular, models were often tuned using data from a very limited set of experimental conditions, and there is typically very little information about how these models work outside the scope of the study they were developed for [1]. Models developed in this way can be severely underconstrained, meaning that many different combinations of the unknown parameters could fit the limited set of target data equally well, so that the parameters that were ultimately chosen may not be realistic, leading to incorrect behavior under different conditions. In many cases, model parameters were tuned manually, and when models were modified to capture new data, the effects of new parameter configurations were typically not assessed systematically. This limits the re-usability of the models and may lead to unnecessary replication of effort.

More recently, several tools have been developed that allow the automated determination of model parameters using a potentially large set of electrophysiological features [14] [15] and support the systematic validation of the resulting models against experimental data [16]. Using the HippoUnit model validation tool, Sáray et al. [16] compared the behavior of several rat CA1 pyramidal cell (PC) models available in the ModelDB database against data from various experimental paradigms. Such comparisons help researchers determine which existing model provides the best fit to the data in the domains they care about and whether they can reuse existing models or should build their own. However, the overall conclusion of this study was that although several models successfully capture certain aspects of the electrophysiological behavior of CA1 PCs (typically those that the models were originally built to replicate), none of the existing models can simultaneously capture all the relevant behaviors of CA1 PCs covered by HippoUnit, and a thoroughly validated, accurate general-purpose model of this cell type is currently missing.

In the current work, we developed and validated morphologically and biophysically detailed models of CA1 pyramidal cells using our previously developed tools for parameter optimization and model validation. We aimed to provide models that are consistent with the widest possible range of experimental observations and are therefore potentially useful for modeling studies with many different purposes. Since this cell type is one of the most well studied to this day, with a tremendous amount of experimental data about its physiological behavior (including both somatic and dendritic behavior under different circumstances such as current injections and synaptic inputs) and its building components (including experimental data about the ion channel distribution and kinetics) that can constrain a wide variety of parameters, it was a solid base to develop a detailed model fitting the aforementioned criteria of wide usability in different studies and to showcase our methods that can be further applied to other cell types.

One crucial aspect of creating an up-to-date biophysical model of CA1 PCs involved reviewing and updating the set of ion channel models used in light of new scientific discoveries. The ability to assess the molecular composition of different ion channels and determine the molecules responsible for ionic currents has improved greatly in the last decade, and the amounts of these molecules can be quantified in the cell membrane, giving computational neuroscientists new quantitative data about channel distributions that can be incorporated into the models, serving as constraints during the parameter fitting process. There is also much more experimental data available about the ion channel kinetics in the pyramidal neurons that can differ in the different dendritic regions. This led to a new curated set of ion channel models specifically for CA1 pyramidal cells, which played a significant role in the improved performance of our model.

To create accurate models of CA1 pyramidal neurons, one also needs to consider the fact that the dendrites of cortical pyramidal cells bear spines that receive most of the excitatory synaptic input, act as separate electrical and biochemical compartments, and play important roles in signal integration and plasticity. Many previous models of CA1 pyramidal cells ignored spines or modeled them in an extremely simplified manner. In this study, we aimed to develop a fully active model including spines to analyze the contributions of nonlinear processes in spines and dendrites to signal integration and action potential generation.

One limiting factor for model development is that modelling dendritic spines in detail greatly increases the computational complexity of the model. Fortunately, in recent years, several modeling tools have been developed that support parallel execution of simulations and can therefore take advantage of the powerful clusters and supercomputers that are now available for scientific research, allowing the efficient creation and execution of complex neuronal models. Nevertheless, we also investigated ways to reduce the computational complexity of models of spiny neurons without altering their functional properties. More specifically, we built and examined simplified versions of our models that did not explicitly model dendritic spines but adjusted membrane properties (including the densities of voltage-gated conductances) using a surface-correction factor (F-factor) that takes into account the membrane area of the spines. Utilizing these tools enabled us to compare different methods for including dendritic spines in our cell models, offering insights about which aspects of neuronal physiology depend on the compartmentalization of specific mechanisms in spines, which also allows researchers to choose the best option for modeling spines in their future studies.

## Results

### Overview of the model-building process

The primary goal of this study was to develop a detailed general-purpose model of rat CA1 pyramidal neurons that properly integrates the current evidence regarding the relevant biophysical mechanisms and successfully captures the electrophysiological behavior of this cell type in a wide range of experimental conditions. To this end, we first systematically processed the available literature on CA1 pyramidal cells (PCs), with particular attention to the biophysical properties and spatial distributions of voltage-gated ion channels (see Methods). This allowed us to identify those parameters (mainly the absolute densities of the various channels in specific cellular compartments) that are not sufficiently constrained by the available data but are expected to have a significant effect on the electrical behavior of the neuron. We then used automated parameter search methods [14] to set these parameters in a morphologically and biophysically detailed model neuron such that the electrophysiological behavior of the model best matched a set of experimental recordings under the same conditions. Finally, we validated the resulting model by quantitatively comparing its behavior to that of real CA1 PCs under a variety of different conditions using data that had not been used during parameter optimization. Figure 1A shows the model-building process.

**Figure 1:**
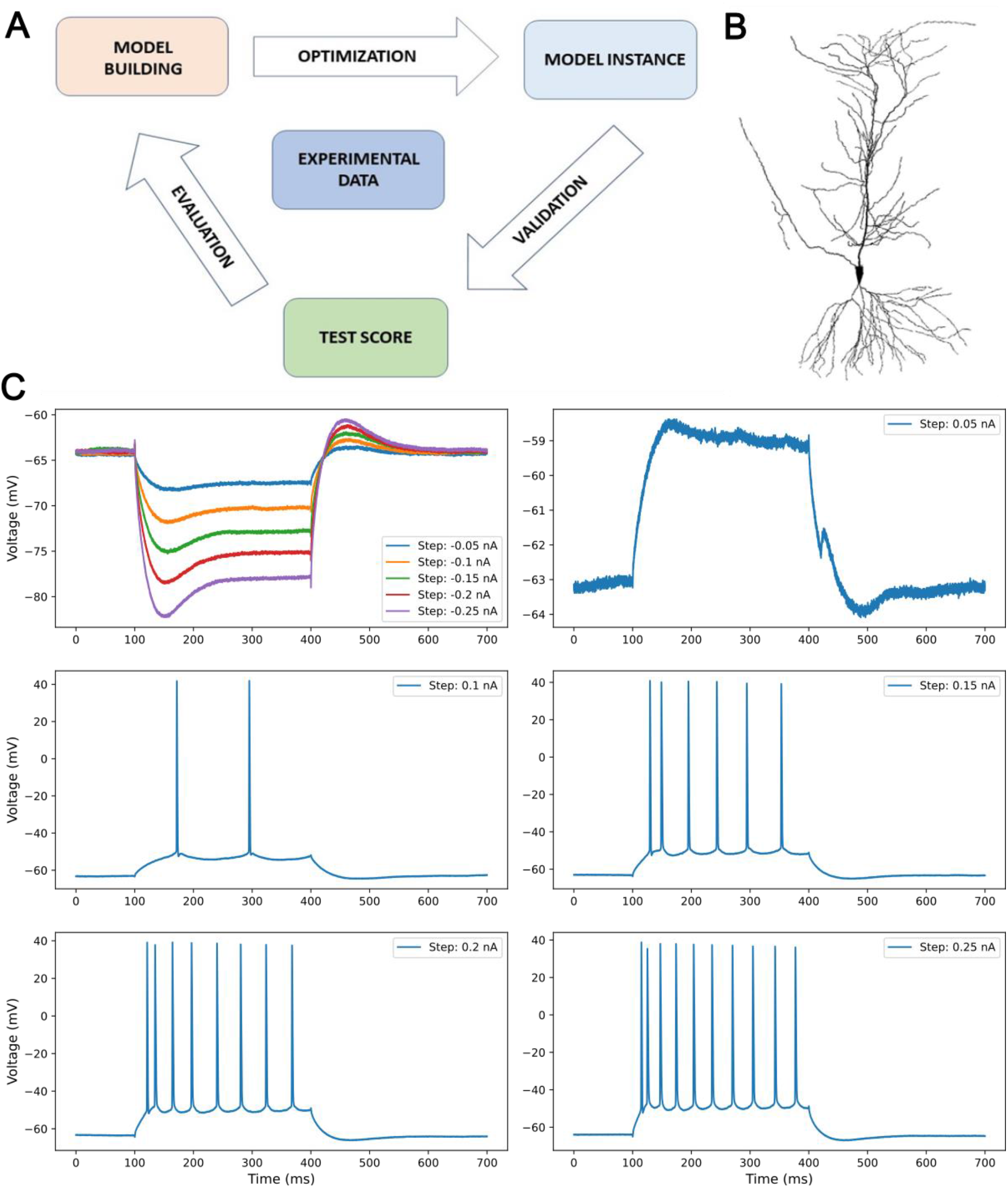
A: A flow chart of our optimization process. First, we build an initial model, then optimize its parameters, which gives back 20 model instances. Then we validate these models with the HippoUnit framework, which gives us test scores. We evaluate our results based on the scores and figures, modify, or expand the models accordingly, and then repeat the entire process. B: The morphology of the model from Megías et al. 2001. while the axon is coming from Bianchi et al 2012. C: An example of the current step protocol and the resulting experimental traces that were the basis of the electrophysiological feature extraction. We extracted features and then optimized our parameters for 5 negative current steps and 5 positive current steps that lasted for 300 ms.

Models were fitted to electrophysiological features extracted from somatic current-clamp recordings from rat CA1 pyramidal neurons (data courtesy of Judit Makara) using the eFEL feature extraction library (https://github.com/BlueBrain/eFEL). Only a subset of experimental features was used for optimization (see Table 2), and the rest were used only for validation to see how the model performs when we evaluate features that were not fitted.

**Table 1:** Each row contains the results of a blocked channel model. The second column contains the expected effect of the blocking based on the experimental literature, while the third and fourth columns contain the effect on the model’s soma and dendrites, respectively. First, the Somatic Features Test was re-evaluated, and then the Backpropagating action potential test. In those cases where the fourth column regarding the dendritic behavior was left empty, there was either no effect on the dendritic behavior, or the somatic behavior changed so much that the individual effect on the dendrites was inseparable.

| Blocked channel | Expected effect | Measured effect |  |
| --- | --- | --- | --- |
|  |  | Soma | Dendrites |
| Sodium channels | No firing | No firing | No firing |
| Dendritic delayed-rectifier | impaired repolarization after spike<br>[34] [26] | No visible effect | bAP: slight change<br>in spike widths on<br>dendrites |
| somatic and axonal delayed-rectifier | impaired repolarization after spike<br>[34] [26], negative shift in voltage<br>threshold for action potentials (APs)<br>and reduced rheobase [71] increased<br>initial firing frequency, depressed<br>maintained firing because neurons<br>entered depolarization block more<br>readily [72] | mini burst |  |
| All delayed-rectifier | impaired repolarization after spike<br>[34] [26] | mini burst |  |
| KD current on axon | effect on the delay in the onset of<br>firing [30] modulation of ADP and<br>bursting [73] | repetitive firing to very<br>small positive currents,<br>elimination of delay |  |
| Ih current | No sag, increased EPSP amplitude<br>and duration [22] | elimination of sag at<br>hyperpolarizing currents | plateau broadening<br>on pathway test |
| All KM current | Destabilized axon, spontaneous<br>firing [74] | spontaneous firing after<br>stim, repetitive firing for<br>small positive currents |  |
| All KA current | Stronger bAP, bigger EPSP [34] [75]<br>[76] [76] | strange firing pattern | no attenuation of<br>bAP |
| KD current on soma | effect on the delay in the onset of<br>firing [30] modulation of ADP and<br>bursting [73] | no visible effect (slightly<br>broader spikes) |  |
| All KD current | effect on the delay in the onset of<br>firing [30] modulation of ADP and<br>bursting [73] | repetitive firing to very<br>small positive currents,<br>elimination of delay |  |
| N-type calcium channel | no ADP and burst firing [73]<br>expecting similar effect as BK<br>channel blocking | firing doublets, strange<br>spike shape |  |
| BK type calc. Dep. Potassium<br>channel | Effect on fast AHP and repolarization<br>of spikes and plateaus [44] | firing doublets, strange<br>spike shape | very broad spikes<br>on soma as the<br>result of synaptic<br>input |
| L-type calcium current | no activation of slow AHP, role in<br>synaptic integration [73] | AHP shape changes |  |
| Slow AHP current | mediating slow AHP | Slight change in the<br>spiking pattern and AHP |  |
| T-type calcium current | impact on pacemaking and repetitive<br>firing [73] | less action potential,<br>visible at 0.15 nA and 0.2<br>nA |  |
| SK type calc. Dep. Potassium<br>channel | regulating medium AHP [77] regulate<br>spike timing precision [78], | more spike at 0.1 and<br>0.15 nA |  |
| R-type calcium current | impact on burst firing [73] | no visible effect |  |

**Table 2:** The optimized eFEL features during the parameter fitting process with the current steps where they were evaluated.

| Feature name from eFEL | Feature description | Fitted current steps (nA) |
| --- | --- | --- |
| Spikecount | Number of spikes during stimulus | -0.05, -0.1, -0.15, -0.2, -0.25, 0.05, 0.1, 0.15, 0.2, 0.25 |
| inv_first_ISI | Reciprocal of first interspike interval | 0.15, 0.2, 0.25 |
| inv_second_ISI | Reciprocal of second interspike interval | 0.2, 0.25 |
| inv_third_ISI | Reciprocal of third interspike interval | 0.2, 0.25 |
| inv_fourth_ISI | Reciprocal of fourth interspike interval | 0.2, 0.25 |
| inv_fifth_ISI | Reciprocal of fifth interspike interval | 0.2, 0.25 |
| inv_last_ISI | Reciprocal of last interspike interval | 0.15, 0.2, 0.25 |
| inv_time_to_first_spike | Reciprocal of time to first spike | 0.15, 0.2, 0.25 |
| steady_state_voltage | Average voltage after stimulus | -0.05, -0.1, -0.15, -0.2, -0.25, 0.05, 0.1, 0.15, 0.2, 0.25 |
| voltage_base | Average voltage during the last 10% of time before the stimulus | -0.05, -0.1, -0.15, -0.2, -0.25, 0.05, 0.1, 0.15, 0.2, 0.25 |
| voltage_deflection | Difference between the voltage base and the steady-state voltage at the end of the stimulus | -0.05, -0.1, -0.15, -0.2, -0.25 |
| voltage_deflection_begin | Difference between the voltage base and the mean voltage in the early phase (5 to 15% of duration) of the stimulus | -0.05, -0.1, -0.15, -0.2, -0.25 |
| AHP_depth_abs | Absolute value of the depth of afterhyperpolarization | 0.15, 0.2, 0.25 |
| AHP_time_from_peak | Time from the peak of the AP to the minimum of the afterhyperpolarization | 0.15, 0.2, 0.25 |
| AP2_amp | Amplitude of second AP | 0.15, 0.2, 0.25 |
| AP_amplitude | Average height of APs | 0.15, 0.2, 0.25 |
| AP_begin_voltage | Threshold of AP initiation | 0.15, 0.2, 0.25 |
| AP_duration_half_width | Half-width of the AP | 0.15, 0.2, 0.25 |
| AP_fall_time | Time from the peak of the AP to the end of the AP | 0.15, 0.2, 0.25 |
| AP_rise_time | Time from the AP threshold to the peak of the AP | 0.15, 0.2, 0.25 |
| AP_last_amp | Amplitude of the last AP | 0.15, 0.2, 0.25 |
| time_to_last_spike | Time to the last spike | 0.15, 0.2, 0.25 |
| sag_amplitude | Difference between the minimal voltage and the steady state voltage at the end of the stimulus | -0.25 |
| sag_ratio1 | Ratio between sag amplitude and maximal hyperpolarization from voltage base | -0.25 |

In every iteration of model fitting, we ran 20 optimizations simultaneously from different, random starting points to avoid solutions where the parameter search process gets stuck in a local minimum instead of reaching the global best solution. Multiple runs also helped us identify when a parameter is consistently tuned near one of its boundaries, which indicated that a better solution could be achieved by extending the relevant boundary. In these cases, the parameter search was re-run with revised boundaries.

During parameter optimization, we used the original reconstructed morphology that did not include spines, and modeled the effects of spines by merging all the relevant biophysical mechanisms (such as the membrane capacitance and various conductances) with the ones present in the dendritic shaft (for details, see Methods and the section “The effect of explicitly modelling dendritic spines on the behavior of the model” below). The dendritic conductance parameters fitted during optimization correspond to their hypothetical values in the equivalent fully detailed model (with spines), but the actual values in the optimized model were calculated by also considering the density of the channel in the spine and the shaft, as well as the F-factor, and simulations were run using this model. Therefore, the maximal conductance parameters seen in Figure 2 are the uncorrected (hypothetical) values, and the maximal conductance values in the simulated spineless model can be different when a particular conductance is present both in the dendritic spines and the dendritic shaft.

**Figure 2:**
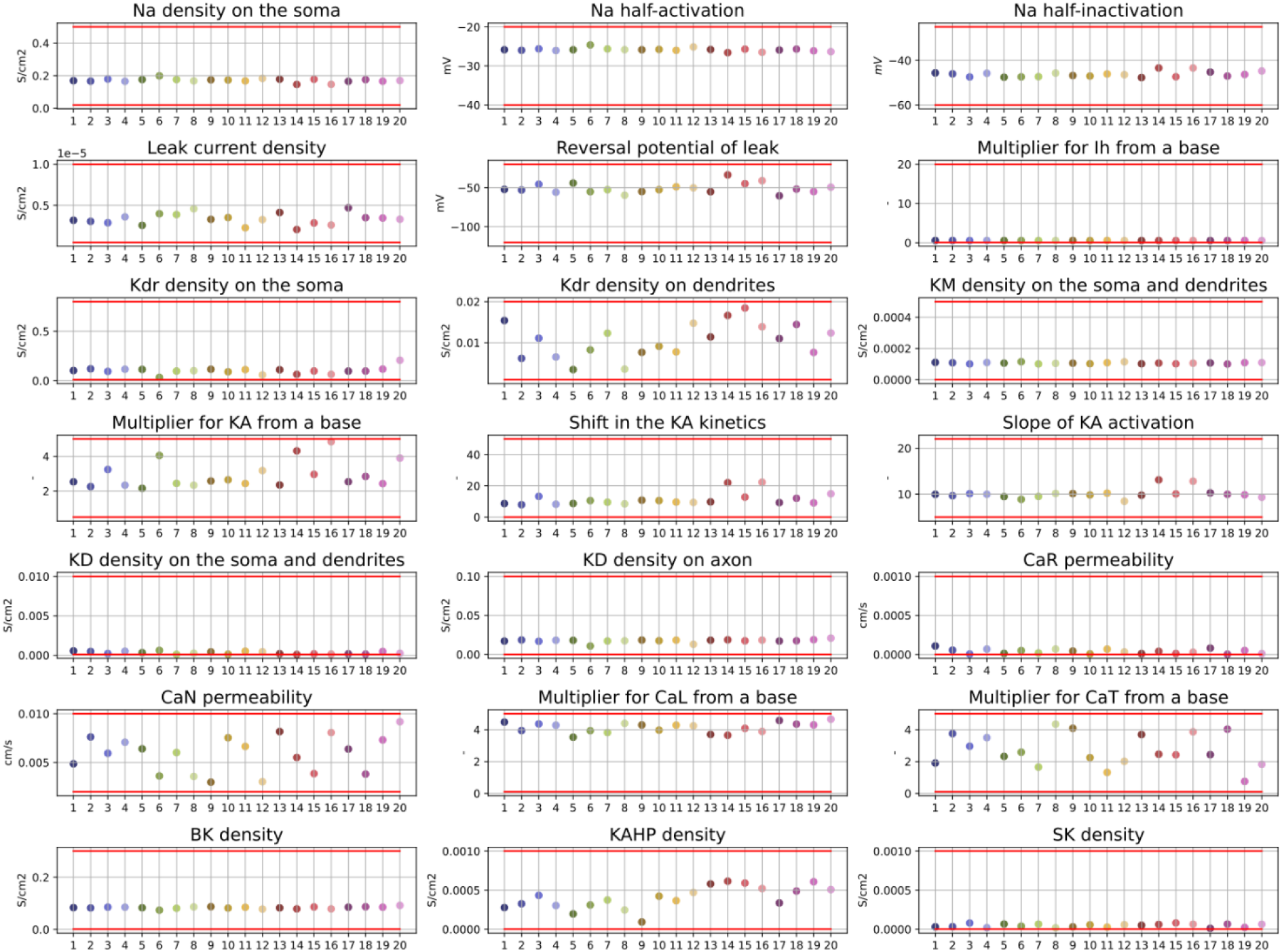
The fitted parameters of the 20 optimized models. The red lines are the upper and lower boundaries of the parameters. The x-axis and the different colors represent the individual models by their numbers, while on the y-axis, there are the parameter values for each model and parameter. Supplementary Figure 1 shows versions of the plots in the panels corresponding to the “Multiplier Ih from base”, “Kdr density on the soma”, “KD density on the soma and dendrites”, “CaR permeability”, and “SK density” parameters using a different scale for the y-axis to allow a better view of the actual parameter values.

To validate the results, we used the HippoUnit software tool, which was developed specifically to compare various electrophysiological properties of detailed models of CA1 PCs with the corresponding experimental data [16]. HippoUnit has five built-in tests. The Somatic Features Test examines the somatic responses of the model cell to step current injections and compares these to experimental results based on the features of the eFEL library; the Oblique Integration Test examines signal integration in oblique dendrites based on the experimental protocol and results of Losonczy and Magee [17]; the Post-synaptic Potential (PSP) Attenuation Test evaluates changes in the amplitude of post-synaptic potentials as they propagate towards the soma [18]; the Back-propagating Action Potential (bAP) Test examines the properties of the action potentials that propagate back from the soma into the dendrites [19]; and the Depolarization Block Test, based on Bianchi et al., 2012 [20], tests whether the neuron goes into depolarization block in response to a sufficiently large somatic current injection and, if so, compares the characteristics of depolarization block with the relevant experimental results.

For candidate models from the parameter optimization phase, we first ran the Somatic Features Test. The outputs of this test include the feature values and feature errors extracted from the models, and the total error score, which is the average of the individual feature errors. We found that all the errors were reasonably small (the vast majority of the feature errors were within 2 standard deviations of the experimental mean), so we moved to other tests. We then evaluated the model from the perspective of the PSP Test and the Depolarization Block Test, then the bAP Test, and then moved to the Oblique Integration Test. Detailed results from these tests are presented below.

The final iteration of parameter search resulted in a set of 20 optimized models. The algorithm found very similar results in the case of parameters that play a role in shaping the somatic responses of the cell, while in the case of parameters mostly responsible for dendritic behavior, we found larger variability since we did not optimize for dendritic features (Figure 2). The models, the full optimization, and all the validation results, along with the scripts for further evaluation, can be found at: https://zenodo.org/records/18907955 and are also available on ModelDB: https://modeldb.science/2031080.

The following sections describe in detail the properties and behavior of these final models, including the construction of the set of ion channel models and synaptic receptor models, the results of a large set of validations, and a comparison of models in which dendritic spines were modeled at varying levels of detail.

### Models of voltage-gated ion channels

The properties, total quantities, and spatial distributions of voltage-gated ion channels are known to be critical for the construction of biologically faithful compartmental models of neurons. Therefore, we carefully evaluated the relevant experimental evidence regarding the voltage-gated ion channels of CA1 PCs and screened the modeling literature for potentially suitable existing channel models. All the candidate channel models were carefully re-evaluated and compared to the latest available data in the literature. The existing channel parameters were adjusted where needed, and if the exact value for a parameter was not available in the literature, it was either fitted by Neuroptimus or, when some kind of relationship was known, was bound to another optimized parameter. This new set of ion channel models greatly improved the model’s ability to reproduce a wide variety of experimental results, since their kinetics were often the source of deviations from the expected behavior. The final version of the model contains the following currents (see Methods for more details).

- Especially in the subthreshold voltage range, the responses of the neuron are strongly influenced by the voltage-independent leak conductance. Since the observed leak conductance is likely the result of multiple ion channels with approximately linear current-voltage relationships but different ion selectivity, we chose to include both the reversal potential and the density (which was assumed to be uniform) of the leak channel among the unknown (fitted) parameters.
- The hyperpolarization-activated current (Ih) mediated by HCN channels also primarily shapes the subthreshold behavior of CA1 PCs. We used one Ih kinetics for the soma and the proximal dendritic arbor and a modified kinetics for the distal dendritic arbor [21]. The amount of Ih increases in the dendrites as a function of distance from the soma [22] [23], so we fixed the proportion of Ih at different distances and fitted only a single scaling factor.
- The model has 3 types of sodium channels with different kinetics for the soma, axon, and dendrites to address the different contributions of Nav1.6 and Nav1.2 channels in the different regions of the cells. [24] The somatic sodium channel density was optimized, while the axonal density was fixed to 40 times the somatic density. On the oblique and basal dendrites, it was 2/3 of the somatic density, and on the tuft dendrites, it was 1/3 of the somatic density based on the data of Lőrincz and Nusser 2010 [25]. On the apical trunk, the density decreased from the somatic value with increasing distance from the soma.

In regard to potassium channels, the model contains delayed-rectifier, D-type, M-type, and A-type voltage-gated potassium channels as well as several calcium-dependent (and, in some cases, also voltage-dependent) potassium channels.

- The delayed-rectifier potassium channel (Kdr) is a well-studied channel [26] [27] [28]. Its main role is to rapidly repolarize the action potential and enable repetitive firing. Delayed rectifier potassium channels are present in every region of the CA1 PCs. We optimized the maximal conductance of the Kdr channels separately on the soma and the dendrites, while on the axon, the amount of Kdr was 40 times higher than the somatic value to match the increased maximal conductance of the sodium channels.
- The D-type potassium current is present in the soma, dendrites [29], and the axon, and its kinetics were carefully examined and adjusted to the literature [30] [31], to get the delay at the beginning of the current step protocol to the first action potential, which can be observed in the experimental data. [30] Since D-type potassium channels are heteromers regarding their α-subunit, which, as homomers, act like delayed-rectifier potassium channels, we found it logical to include D-type potassium channels in the axon and the soma as well, but their maximal conductance was separately fitted as parameters due to the lack of information. [26]
- M-type potassium channels are present on the soma, axon, and dendrites. The kinetics came from Hönigsperger et al. [32] (see Methods for details). The maximal conductance was the same for the soma and dendrites, while on the axon, the maximal conductance was 4 times the somatodendritic value. [33]
- A-type potassium channels are present in the soma and the dendrites, and their maximal conductance increases with the distance from the soma. [34] We fitted a multiplier parameter that changes the amount of A-type potassium current proportionally on the whole dendritic arbor from a starting point that is based on Kerti et al. 2011. [35] We also introduced a shift parameter that could modify the voltage dependence of the activation during our optimization process. It is known that A-type potassium channels can undergo different kinds of modulations that alter the channel kinetics [36] [37] [38] [39] [40] [41], which gives us some freedom to modify the kinetics to better fit the model’s needs.
- The model contains N-type calcium channels exclusively on the soma. The modeling of the N-type calcium channel follows the Goldman-Hodgkin-Katz equation, with its permeability being a fitted parameter. Its kinetics are based on McNaughton et al. 1997. [42]
- High-voltage-activated L-type calcium channels play a role in shaping the currents in the dendrites and spines of the CA1 pyramidal cell. It is also important to note that calcium ions entering through the L-type calcium channels are responsible for activating the calcium-dependent potassium channel that shapes the slow afterhyperpolarization. [43] L-type calcium channels are present on the soma and dendrites, but the amount of the channels is linearly decreasing with the distance from the soma. [44] [45] The channel kinetics are based on Bell et al. 2001. [46]
- The T-type calcium channels kinetics were based on Randall and Tsien 1997. [47] while the approximate distribution in each dendritic region came from McKay et al. 2006. [48] Since the densities measured by McKay et al. did not seem to follow any simple rule, the relative amount of T-type calcium channels in the model was set directly according to their data (see Methods). Since McKay et al. measured the three different subtypes of the T-type calcium channels separately, in this study, the summed amount was used for our channel model as a baseline amount, and then we used a multiplier as a fitted parameter so the ratios of the maximal conductances between the dendritic regions were preserved.
- R-type calcium channels have a major role in shaping the EPSP and EPSC in the spine heads during synaptic stimulation. [49] One spine head can contain 5-15 channels [50]; based on this and the known single-channel conductance of Cav2.3 (the subunit responsible for the R-type current in the CA1 pyramidal cells), we could calculate the feasible range of permeability values to set the boundaries for this optimized parameter. The properties of the channel were described by Foehring et al. 2000. [51] and Randall & Tsien 1997 [47]. Since the optimized models do not have explicitly modeled spines, which are instead taken into account by the increased membrane area of the dendrites, in these models, the R-type calcium channels are effectively present on the dendritic shaft, which may artificially increase their role in shaping the cell’s response to different inputs. We will examine the effects of modeling spines at different levels of detail in a later section.
- Big-conductance (BK)-type calcium and voltage-dependent potassium channels read out the calcium concentration exclusively from the N-type calcium channels, since according to the literature, these two kinds of channels are only a few nanometers apart from each other in the membrane. [44] The BK-type potassium channel has a role in shaping the fast afterhyperpolarization and spike repolarization [52]. The BK-type potassium channel that works best in our cell model originates from Maurice et al. 2004. [53] It has both voltage and calcium-dependent activation modeled as two separate gates since we have no information whether the voltage influences the calcium dependency of the activation.
- We also introduced the Small-conductance (SK) calcium-dependent potassium channel to the model. It is present in the soma, the dendrites, and spine heads (2-3 channels/spine). According to Ballesteros et al. 2012. [54] the number of SK-channels increases linearly with the distance from the soma. The somatic value was a free parameter between boundaries close to the experimental data [54], and from that, the number of SK-channels increases with distance from the soma. The channel kinetics were based on Hirschberg et al. 1998. [55].
- Connected to the L-type calcium channels, we introduced a generic calcium-dependent potassium channel to the model that is responsible for slow afterhyperpolarization, since there is contradictory information about the identity of this channel. [56] Its kinetics were based on Traub et al. 1991. [57], it was only present on the soma, and the maximal conductance of the channel was a fitted parameter.

Dendritic spines are specialized compartments of neurons, both in a biophysical and biochemical sense. They play critical roles in the generation of excitatory postsynaptic potentials and in synaptic plasticity. Therefore, we paid special attention to identifying the biophysical mechanisms (voltage-gated and ligand-gated ion channels) that shape electrical responses in dendritic spines. Based on the literature, we included the following voltage-gated channels in the dendritic spines of our model neurons.

- A-type potassium channels, with their maximal conductance density matching that of the parent dendrite from which the spine originated. Depending on the location of the spine, either proximal or distal A-type potassium channels were used.
- Small-conductance (SK-type) calcium-activated potassium channels, whose maximal conductance density also matched that of the parent dendrite.
- L-type calcium channels, whose density (permeability) also came directly from the originating dendritic location.
- R-type calcium channels, whose permeability was determined by the parameter fitting process.
- T-type calcium channels, whose permeability was provided directly by the experimental data [48] and depended on the dendritic region where the spine was located, except for the trunk, where the permeability of the T-type calcium channel equals the number on the shaft in the absence of more accurate data. The permeability on the dendritic shaft is a fitted parameter.

### Model validation

In this section we describe the behavior of the optimized models mainly through the application of the HippoUnit tool, which was developed specifically to enable the comprehensive, quantitative validation of detailed models of CA1 PCs [16]. However, there are a few basic properties of these neurons that are not currently included in the HippoUnit framework (and that were not directly optimized for during the parameter search), which are nevertheless important indicators of the physiological behavior of CA1 PCs. These properties, which were evaluated separately before the application of HippoUnit, were the following:

The cells should not fire spontaneously before current injections, and the model should stop firing when the current injection ends. It was also a requirement for the spikes to initiate in the axon (Supplementary Figure 2). Finally, it was important that our model should not enter depolarization block in response to the optimized current injections.

Supplementary Table 1 contains the final scores of the models on each HippoUnit test. All 20 optimized models gave very similar results, which proves that Neuroptimus could find the global minimum error from random starting locations.

#### Somatic Features Test

We first evaluated the models on the somatic features test. The resulting traces and the wider variety of features allowed us to get a more in-depth understanding of the models. In Figure 3, there is a selection of representative and diverse features of the models, and the models’ results compared to the experimental data. Note that this is only a selection of our choice for demonstrative purposes. The whole validation results can be found at: https://zenodo.org/records/18907955. A typical model’s voltage response to the somatic current injection protocol used in this test can be seen in Supplementary Figure 3.

**Figure 3:**
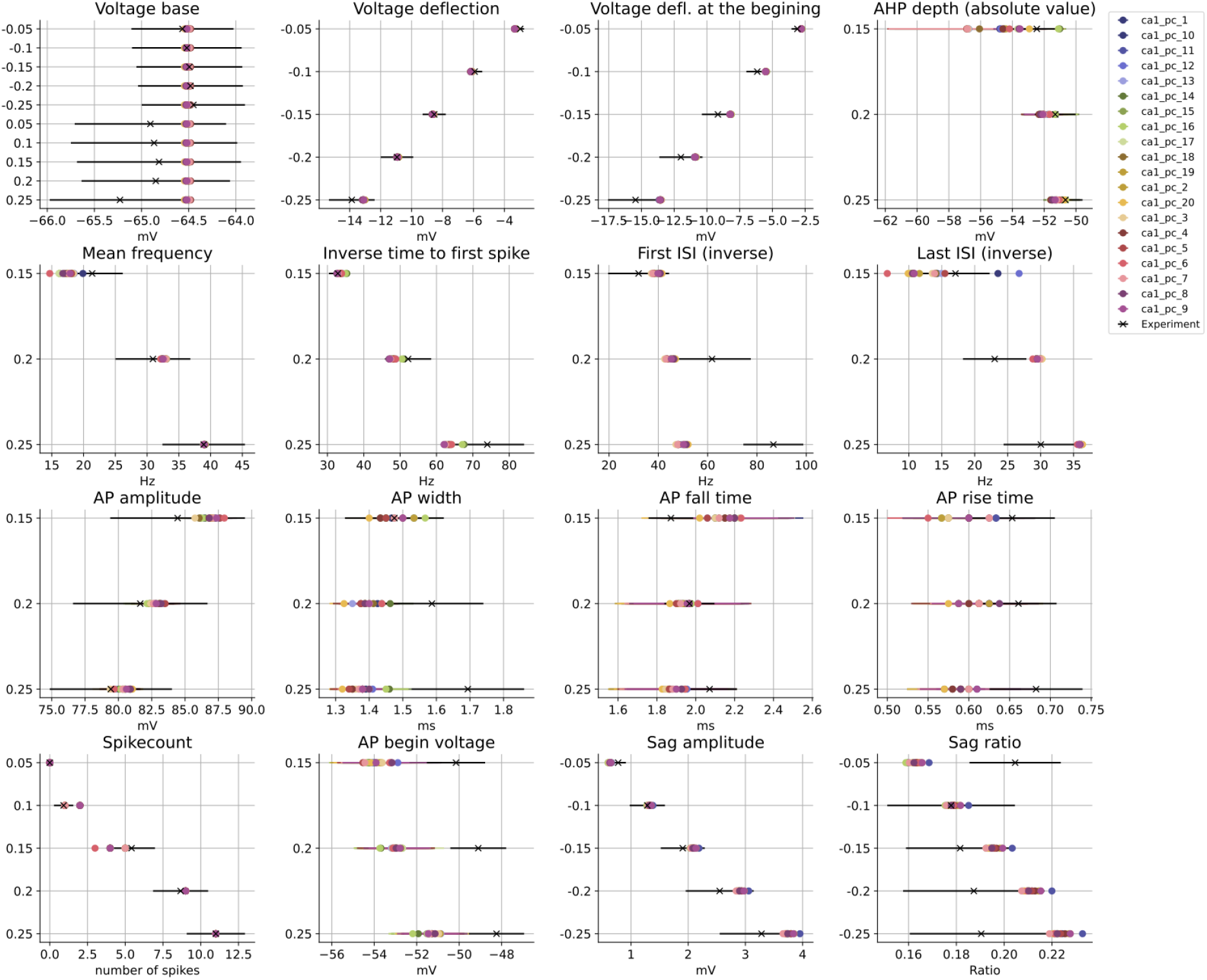
The result of the Somatic Features Test of the HippoUnit for the 20 optimized models. For this figure, 16 representative features were selected to show the wide variety of the somatic behavior of the cells, from passive properties to action potential shapes and spiking patterns.

#### Backpropagating Action Potential Test

Getting backpropagating action potentials right was crucial for the further examination of the dendritic integration of the models. The amplitude of the backpropagating action potential depended on the amount of sodium channels, and the difference between the first and the last action potential amplitude depended on the inactivation of the sodium channels on the dendrites (see Supplementary Figure 4). All of the models were “strong-propagating” type of cells based on the experimental observations [19]. Identifying the necessary conditions for weak propagation was outside of the scope of this study, and this issue was previously investigated by Bernard and Johnston (2003.) [58] [19]. Results of the Backpropagating action potential test are in Figure 4A,B.

**Figure 4:**
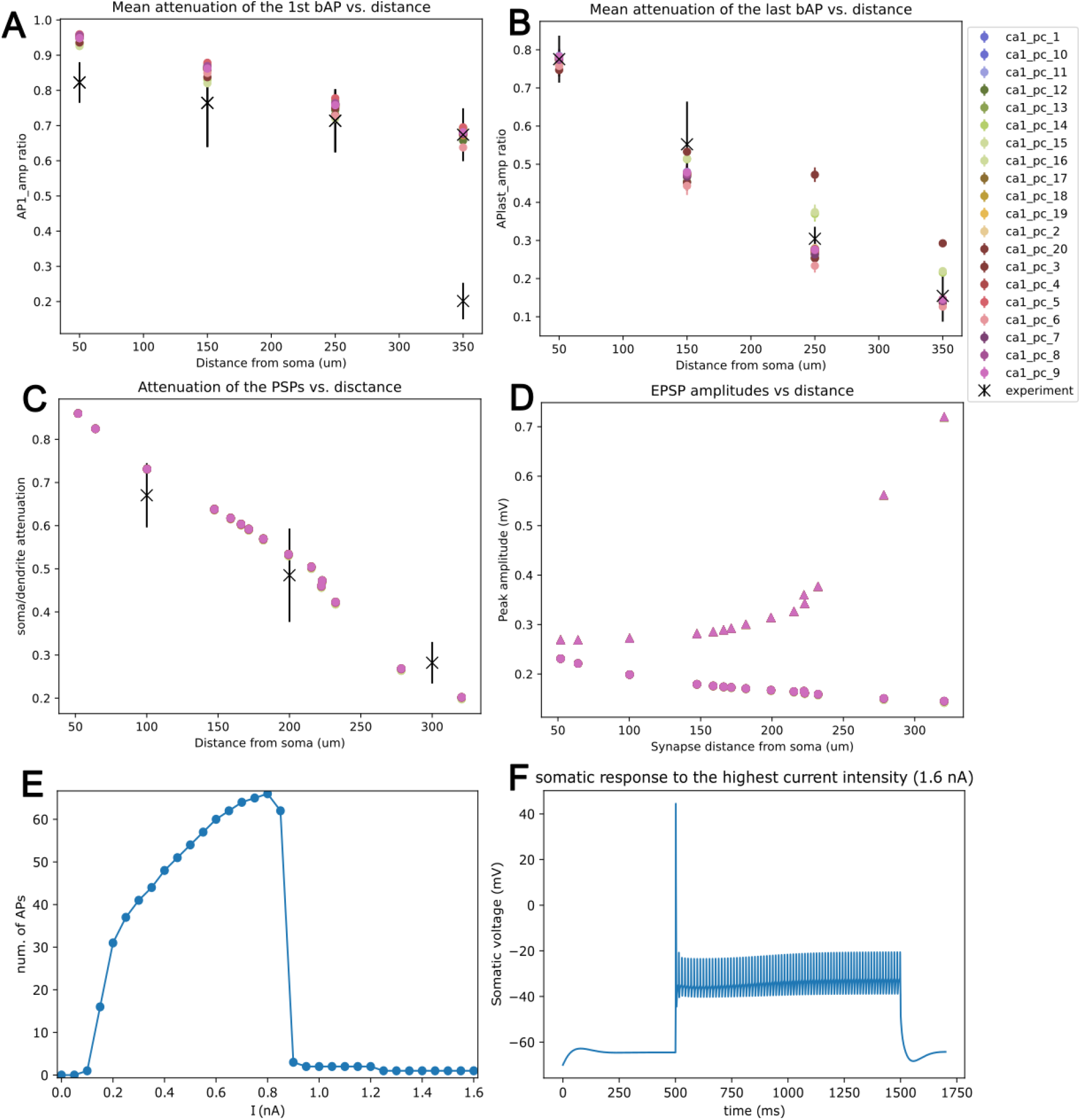
Validation results of the 20 optimized models. A shows the first action potential propagating from the soma on the apical trunk to the distal dendrites and B shows the last action potential of the model propagating from the soma on the apical trunk to the distal dendrites that are results from the Back-propagating action potential Test of the HippoUnit, that shows the first and last action potentials propagating from the soma on the apical trunk to the distal dendrites. The models show very little variability. C and D show the result of the Post-synaptic potential attenuation test, where an EPSP propagates towards the soma. The models show no variability, because the result of this test only depended on the axial resistance, which was the same in every model. E and F show an example of the results from the Depolarization Block test. E shows the FI curve of a model, which could be interpreted as a depolarization block, but the actual trace on F shows that there are spikes, but they become smaller than the detection threshold of -20 mV.

#### Post-synaptic Potential Attenuation Test

We found that the attenuation of post-synaptic potentials depended mainly on the passive properties of the cell model. Based on the recent in vivo findings about a stronger coupling between the soma and the dendrites [59] and computational observations made in our lab [60], we lowered the axial resistance of the model from 100 to 50 Ohm/cm^2^,which resulted in a good fit to the experimental data (Figure 4C, D). Using the traditional 100 Ohm/cm^2^ axial resistance resulted in too strong attenuation (see Supplementary Figure 5), which remained the same after re-optimizing the model, and was therefore independent of the fitted parameters.

#### Depolarization-block Test

The Depolarization Block Test examines whether the soma enters a depolarization block and how much injected current is needed for that. According to the test, the performance of the models varied: there were some that entered depolarization block, but others did not. However, looking at the individual traces, we concluded that, in fact, all the models showed qualitatively similar behavior: in response to large current injections, the amplitude of somatic APs was reduced drastically, but small spikelets always remained. The result of the test depended on whether these spikelets reached the -20 mV detection threshold. If they did, the test reported that the model did not enter depolarization block, and when they did not, it was considered a depolarization block. An example of this phenomenon can be seen in Figure 4E, which shows the f-I curve of the model, and Figure 4, which shows the corresponding voltage trace. The spikelets themselves originate from the axon, then backpropagate to the soma (Supplementary Figure 6). We conclude that the soma could enter a real depolarization block in principle, but the axon currently prevents this. Despite our model failing this HippoUnit test, there is experimental evidence that supports the presence of spikelets coming from the axon. [61] A more detailed and fine-tuned axon could probably help to capture this phenomenon more accurately.

#### Finding the correct voltage dependence of the NMDA receptor model using the Oblique Integration Test

Cortical pyramidal neurons, including CA1 PCs, implement highly nonlinear forms of dendritic integration, which are mediated by several distinct mechanisms such as Na spikes, Ca spikes, and NMDA spikes or plateaus [62]. More specifically, the thin apical oblique dendrites of CA1 PCs show supralinear summation of synaptic inputs arriving at nearby dendritic spines if these inputs are activated within a short time window, and the phenomenon depends on the activity of both dendritic voltage-gated Na channels and NMDA receptors [17]. The Oblique Integration Test of HippoUnit replicates some of the key experiments in [17] to compare the characteristics of synaptic integration in oblique dendrites of detailed CA1 PC models to the corresponding experimental data [16].

Unlike the previously presented validations that did not utilize synapses or (in the case of the PSP Attenuation Test) used only a standardized AMPA-type synaptic current input, the results of the Oblique Integration Test are strongly influenced by the properties of synaptic inputs, and it was therefore crucial to have a realistic model of excitatory synapses that are activated by the Schaffer Collaterals in this dendritic region. These inputs are known to activate both AMPA- and NMDA-type glutamate receptors, so it was important to properly capture the conductance ratio of the two receptors as well as their kinetics, and especially the voltage dependence of NMDA receptor activation, which is a critical factor that determines the emergence and properties of NMDA spikes/plateaus.

By default, HippoUnit uses the NMDA receptor voltage dependence from the model of Jahr and Stevens [63] and an AMPA/NMDA conductance ratio of 2.0 in the Oblique Integration Test. However, this configuration produced sublinear dendritic integration and a correspondingly high error in the test. On the other hand, the experimental and modeling literature contains several competing models of NMDA receptors with distinct voltage dependence, and there are also relevant experimental results that constrain the AMPA/NMDA ratio in Schaffer collateral synapses. Therefore, we systematically evaluated oblique dendritic integration with several different models of NMDA receptors and recalculated the AMPA/NMDA conductance ratio based on the available experimental evidence. We determined the AMPA/NMDA conductance ratio separately for each NMDA receptor model based on the experimental study of Otmakohova, Otmakhov, and Lisman [64]. These authors used somatic voltage clamp and pharmacological blockade of NMDA receptors to measure the contributions of AMPA and NMDA receptors to the excitatory postsynaptic current (EPSC) evoked by the activation of the Schaffer collaterals or the perforant path and determined the ratio of the total charge carried by AMPA vs. NMDA receptors by calculating the areas under the EPSC curves corresponding to the two types of receptor. We replicated their experiments in silico (see Methods for details) and determined the AMPA/NMDA conductance ratio that was necessary to reproduce the measured total charge ratio.

We used the Oblique Integration Test of HippoUnit (with custom NMDA receptor models and AMPA/NMDA conductance ratios) to analyze how the voltage dependence of NMDA receptor activation shapes the nature of synaptic integration in thin dendrites of CA1 PCs, and to determine the voltage dependence that provides the closest match to the experimental data. First, we evaluated models with eight distinct NMDA receptor voltage dependences from different published studies, including Jahr and Stevens [63], which is the source of the default NMDA receptor model in HippoUnit; Major et al. [65]; Ujfalussy [66], in which voltage dependence was deliberately placed between those of the Jahr-Stevens and Major models; Grunditz et al. [67]; Otmakhova [64]; Larkum [68]; Magee [69]; and Gao [70].

We ran the Oblique Integration Test [16] using each of these voltage dependences (Figure 5B) but otherwise identical synaptic parameters, and compared the resulting features, error scores, and the individual voltage traces. The test first selects candidate dendrites with the required geometric properties, then creates an active dendritic spine at the chosen location, which will receive the synapses. Next, it determines the synaptic weight (which scales the maximal conductance of both AMPA and NMDA receptors) such that a jump in the maximal steepness of the somatic voltage response (dV/dt) occurs as the number of activated synapses is increased from 4 to 5, which was a typical scenario in the experiments of Losonczy et al. [17]. This jump is typically associated with the appearance of a dendritic Na spike in the stimulated dendritic branch. Then it runs the main simulation experiments where increasing numbers of synapses are activated either synchronously or asynchronously on a given dendritic segment, and the somatic and dendritic voltages are recorded. Finally, it calculates various features such as the apparent voltage threshold for supralinear behavior, the amplitude of the response at threshold, and the degree of nonlinearity at and above the threshold.

**Figure 5:**
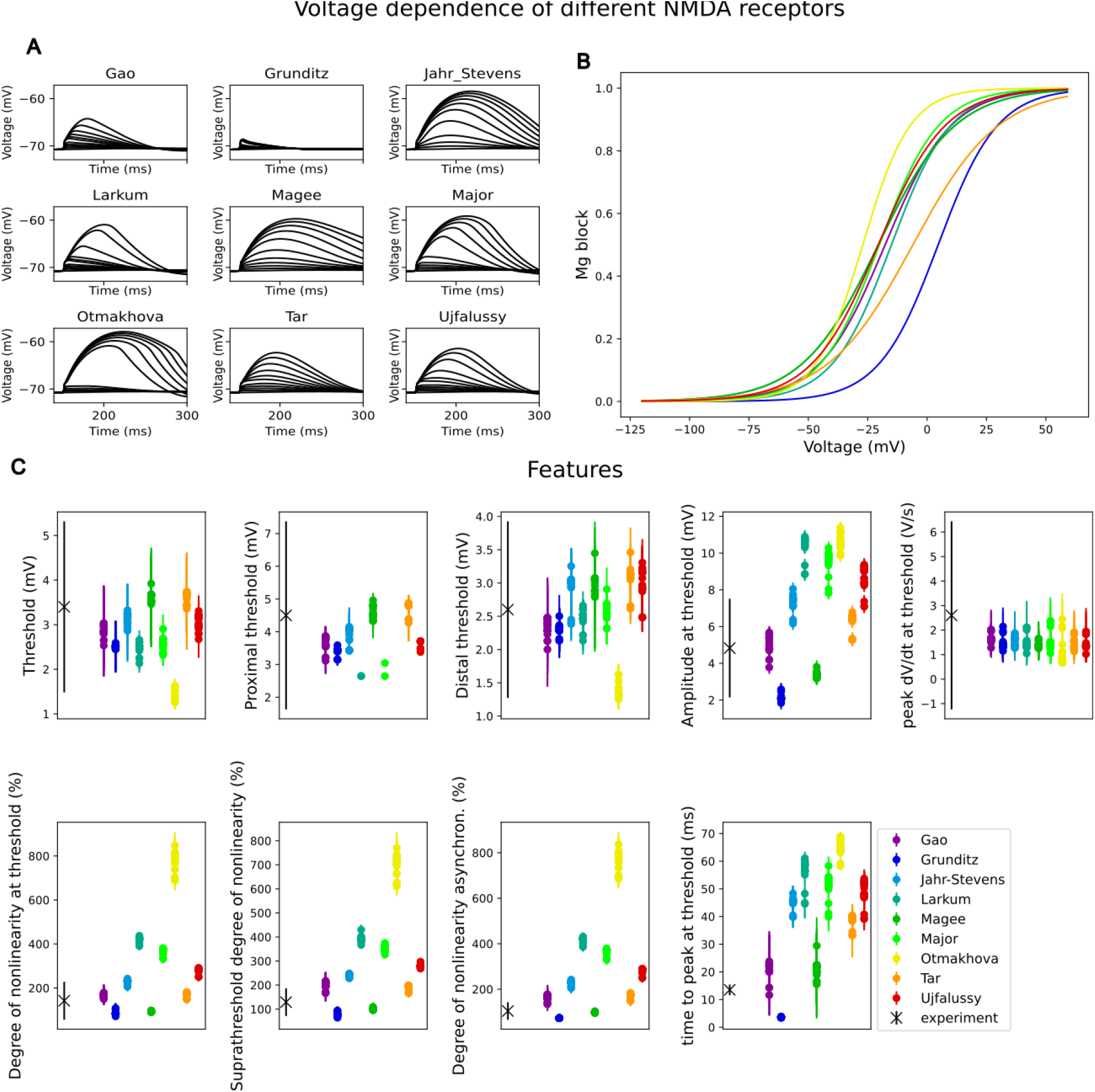
Voltage dependence of different NMDA receptors. A) shows an example of the results of the HippoUnit’s Oblique Integration test protocol, the response to an increasing number of inputs on a distal dendrite measured on the soma with different NMDA receptor models. B) Shows the voltage-dependence of the different NMDA receptor models, and C) shows the features of the Oblique Integration test and how the models behaved in the case of each NMDA receptor.

We found large differences between the results of the Oblique Integration Test between models with different voltage dependences in the NMDA receptor, both at the level of the actual voltage responses to synaptic stimuli (Figure 5A) and in the values of several calculated features (Figure 5C). The voltage dependence of the NMDA receptor had a particularly strong effect on all the features that measure the degree of nonlinearity: some models showed a much greater degree of supralinearity than real neurons, some others remained sublinear, and only in a few cases did the degree of nonlinearity match the experimental results. The EPSP amplitude at threshold was similarly variable. On the other hand, some other features, such as the peak dV/dt at threshold, were not significantly affected by the voltage dependence of the NMDA receptor because this feature reflects the effect of dendritic Na spikes and depends mainly on the amount and properties of dendritic Na channels.

When looking at the somatic voltage responses to varying numbers (1-10) of synchronously activated synapses (Figure 5A), we observed prominent differences in the appearance of the long-lasting, large-amplitude component of the response that reflects non-linear activation of NMDA receptors (NMDA plateaus). In some cases, no NMDA plateau could be seen (Gao, Grunditz); in other cases, the NMDA plateau appeared only at input numbers much larger than the threshold (Larkum, Major, Ujfalussy, Magee); finally, in some cases, there was a very large jump in the response amplitude due to the appearance of the plateau, with little further increase as the number of synapses increased further (Otmakhova). None of these patterns matches the experimental observations of Losonczy et al. [17] and therefore rules out the corresponding NMDA receptor voltage dependence in our model.

Overall, we found that none of the previously proposed voltage dependences led to completely satisfactory results in our model, so we decided to tune the parameters of the sigmoid function to achieve better results. By systematically changing the half-maximal point and the steepness of the sigmoid, we found a voltage dependence that led to oblique integration behavior matching experimental data both qualitatively and quantitatively (according to the scores of the Oblique Integration Test of HippoUnit). We found that the ideal NMDA receptor model for our model cell activates roughly at the same voltages as the receptor model of Ujfalussy et al. but then continues flatter, having a half-activation at -6 mV and at higher voltages it looks more like the NMDA receptor model of Grunditz et al. (Figure 5B) With this voltage dependance the model could replicate the jump driven by a dendritic Na spike at 5 inputs and had the desired degree of nonlinearity while avoiding all of the previously mentioned issues regarding the amplitude of the NMDA component at higher input numbers.

#### The roles of specific ion channels in the model

To verify that the ion channel models have the same effect on the model as they have in a real cell, we systematically blocked each of them and re-evaluated the HippoUnit tests in each case. The detailed results of the channel blocking can be found in Table 1 and Supplementary Figure 7. In summary, each ion channel model that is present in the models is qualitatively responsible for the phenomena that are stated in the literature. Delayed-rectifier is not the only contributor to the action potential repolarization, but blocking it impairs the spiking pattern, while D-type potassium channels play a big role in the excitability of the cells and determine the delay of the first spike in a spike-train. M-type potassium current also plays a role in the excitability of the cell and ensures that the models stop firing when the current injection ends. Blocking A-type potassium channels had a big role in shaping the backpropagating action potentials on the dendrites and disarranged the spiking pattern and the shape of the action potentials. Blocking BK-type potassium channels led to strange action-potential shapes and enabled the cell to fire in doublets. Interestingly, in response to synaptic input, blocking these channels led to a wide action potential on the soma. Because of the strong coupling with the N-type calcium channels, blocking the calcium channels led to the same result.

#### The effect of explicitly modeling dendritic spines on the behavior of the model

All the results that we have presented so far were obtained with models in which most dendritic spines were not explicitly modeled. More specifically, for the parameter optimization and for all validations except the Oblique Integration Test, we modeled all dendritic spines indirectly, essentially merging their membrane surface (including all the spine-associated conductances) into the model of the corresponding dendritic segment (see Methods). For the Oblique Integration Test, we explicitly modeled all the spines that received synaptic input at any stage during the test, but we still modeled the rest of the spines using the F-factor method (modified to exclude the explicitly modeled spines; see Methods). The main reason for using these morphologically simplified models in these in silico experiments was that explicitly modeling tens of thousands of spines increases the computational demands of simulating the model (including the simulation time) by more than an order of magnitude. This becomes a serious issue when many simulations are required, such as during parameter optimization, even with the use of high-performance computing resources. However, to validate our methodology, we carried out an explicit comparison of different versions of our final models with either explicitly or indirectly modeled dendritic spines.

We constructed and simulated three different versions of the models. In the first version, all the spines were modelled indirectly with the F-factor method. This was our baseline model, and this was also how the parameters were fitted during the optimization process. In the second case, only those spines that received synaptic input were explicitly modelled. These models effectively differed from the first case only in tests where there was synaptic input, i.e., the Oblique Integration Test. Our third case was where all dendritic spines were individually modelled, whether they got synaptic input or not. In this case, all the HippoUnit tests were re-evaluated.

In the Somatic Features Test the optimized models and the models where all spines were present behaved very similarly. We found only a slight difference in the resting membrane potential (voltage base), where the models with all the spines were hyperpolarized by 0.5 - 1 mV compared to the optimized model. Every other difference was the indirect result of this hyperpolarization. Its origin can be traced back to the T-type calcium channels that are active at the resting membrane potential, and some of them were relocated to the spine heads in the all-spine case (Supplementary Figure 8).

In the case of the Backpropagating Action Potential Test, the PSP attenuation test, and the Depolarization Block Test, the cells that had all their spines modeled explicitly behaved in exactly the same way as the optimized models (Supplementary Figure 9).

However, more differences could be found between the models in the Oblique Integration Test. We compared three different cases: one where no spines were modelled, and the synapses targeted the dendritic shaft, one where only the spines that received synapses were modeled, and a third where all the dendritic spines were modeled. In the second case, we modified the Oblique Integration Test to first put a spine at the chosen dendritic location before connecting the synapse to the spine head. Test results and comparisons of the optimized models versus the spiny models can be seen in Figure 6. Figure 6B, C, and D show that in the cases when synapses arrive at the spine head, it does not matter if we model all dendritic spines or just the ones receiving synapses. However, the results were clearly different between the cases when synapses arrived on the shaft versus the spine. The degree of nonlinearity at threshold in the case when synapses arrived on the dendritic shaft was lower than the experimental data, but when synapses arrived on the spines, the signal integration became supralinear. We investigated the individual traces to determine the reason for this difference. We found that the sodium spike at the beginning of the voltage response is slightly lower when synapses arrive on the spine heads, but the NMDA component has a higher amplitude and longer duration (Figure 6A), which is responsible for stronger nonlinearities in signal integration.

**Figure 6:**
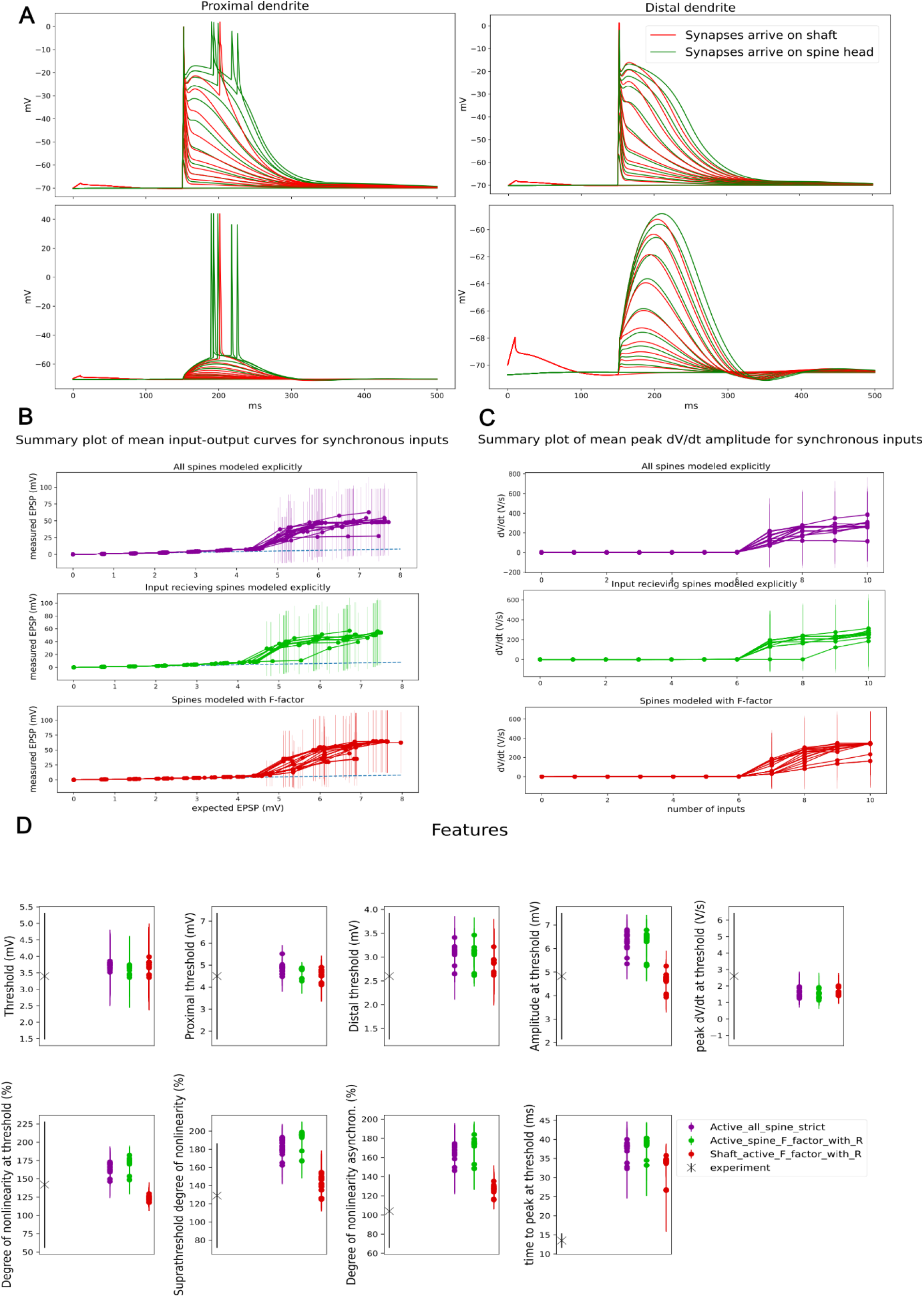
A) An example of the voltage traces resulting from the Oblique Integration test of the HippoUnit. The first row shows the dendritic traces when we put synapses onto the proximal part of the dendrite (first column) or the distal part of the same dendrite (second column), and the second row shows the resulting voltage measured on the soma. Red traces are the result when we put the synapses on the dendritic shaft, and green when we put the synapses on a spine head at the same dendritic location. On B, C, and D, validation results of the Oblique Integration Test in 3 different cases are shown: red: the spines were modeled with the F-factor method, these were our optimized models, green: those spines that received synaptic input were modeled explicitly, the others were taken into account by the F-factor method, purple: all spines were modeled explicitly. B) Shows the mean EPSPs vs. the expected EPSP amplitude in the case of linear summation of the input. Each line is a mean and standard deviation of the different dendrites of a model, and the lines represent models with different parameter sets. C) Summary plot of the peak dV/dt amplitudes in the same fashion as B). D) The absolute values of the features of the Oblique Integration test. The cases are very similar to each other and to the experimental data (except for one feature).

## Discussion

In this paper, we described the systematic and detailed modeling process that allowed us to build a new up-to-date model of CA1 pyramidal neurons that captures a wide variety of different cell behaviors and is expected to be useful in many different modeling scenarios. In addition to *in silico* studies of hippocampal pyramidal cell physiology, the model can be used, among others, as a key component of realistic models of hippocampal network dynamics and function [9] [13], but also as a basis for studying the interactions between biophysical and biochemical processes in various neuronal compartments, including dendritic spines.

One crucial aspect of the new model is its updated and carefully evaluated set of ion channel models, which should be a useful resource in itself for the hippocampal and the broader neural modelling community. Our results also highlight the need for more high-quality, cell-type-specific experimental data on the key properties of ion channels, including their molecular identity, intracellular spatial distribution, and gating via the membrane potential, ionic concentrations, and various biochemical modulators.

Our results also demonstrate the utility of our previously introduced software tools for the automated optimization of model parameters [14]and for the systematic validation of single cell physiological behavior [16], and we hope that the current showcase will encourage the modeling community to include these or other similar tools in their workflows, which in turn would increase the reproducibility, robustness and transparency of computational neuroscience research.

### Modeling dendritic spines

Dendritic spines contribute substantially to the membrane surface of cortical pyramidal cells, receive the majority of excitatory synaptic inputs, and are believed to play important roles in electrical and chemical signaling in these neurons [79] [80] [81] [82]. On the other hand, due to their large number, modeling all spines in detail increases the computational cost of simulating the electrical behavior of pyramidal cells by at least an order of magnitude, even using simplified models of spine morphology. Therefore, many previous studies, starting from the seminal work of Rapp et al. (1992) [83], used models that, rather than modeling each individual spine explicitly, captured their overall effect on dendritic electrical signaling by adjusting the membrane capacitance and membrane resistance of spine-bearing dendritic shafts to account for the surface of the corresponding spines. This approximation has been shown to work well when the spines are passive (i.e., they contain no voltage-dependent mechanisms) and the spine head and the spine base are approximately isopotential, which generally holds in the absence of external (synaptic) input to the spines [84] [85] [86]. Several studies have also successfully applied a hybrid approach where spines receiving synaptic input are modeled explicitly while the rest are taken into account via the F-factor method [87], and we also followed this route in most of our simulations.

However, the spines of CA1 PCs are known to be active, contain several voltage-gated and calcium ion concentration-dependent ion channels, and receive synaptic inputs that are partly mediated by voltage-dependent NMDA receptors. We aimed to capture these mechanisms in our detailed model, and we therefore developed and tested a version of the model where all spines were explicitly modeled, including these nonlinear mechanisms. We compared the behavior of this full model with the simplified version that resulted from parameter optimization, where the spines were omitted but, analogously to the F-factor method, all the conductances (including active and passive ones) as well as the capacitance of the spine were merged into the model of the dendritic shaft. Although some previous studies suggested that this method may lead to incorrect results [88] [89], we found that the two versions of our model gave almost indistinguishable results regarding the somatic voltage response to somatic current injections, the attenuation of single postsynaptic potentials from various dendritic sites to the soma, and even the backpropagation of somatic action potentials into the apical dendrites. On the other hand, the model without any explicitly implemented dendritic spines behaved quite differently from the full model when we looked at the integration of synaptic inputs to oblique dendrites, which was expected since this paradigm is based on synaptic inputs to dendritic spines, and the experimental results, which are successfully replicated by the full model, are known to depend on various voltage-dependent mechanisms including the nonlinear NMDA response at the synapse.

However, even in this case, results very similar to those of the full model could be obtained from a third version of the model where only the spines receiving synaptic input were modeled explicitly. The computational cost of this model is intermediate between the full model and the one without any spines, although the exact cost depends on the number of spines that receive active synapses. Therefore, models of this type can be a good choice when synaptic inputs are important, but the resources available for simulating a single neuron need to be kept reasonably low, which is the typical case for large-scale realistic network models, and also for exploring the parameter space of single neurons responding to realistic patterns of synaptic input. However, it should be noted that such hybrid models can be used only if it is known *a priori* which spines may receive synaptic inputs during the simulation.

### Challenges in automated parameter fitting

Building models systematically and fitting their parameters to experimental data while keeping all the other properties of the chosen cell type that are not included in the parameter fitting process in mind raises several challenges. Our final set of models was created by optimizing neuronal parameters (mostly the densities and, in a few cases, the properties of ion channels) to match the somatic voltage response of real CA1 PCs to somatically injected step currents, as measured by a carefully chosen (sufficiently rich, but not strongly redundant) set of physiological features. Parameter optimization was carried out in a fully automated manner; however, a substantial amount of human expertise was still required to set up the optimization, e.g., to define the mechanisms included in the model, to choose the parameters to be optimized as well as their plausible ranges, and to set the values of the fixed (non-optimized) parameters. Our choices were driven mainly by the available experimental data, which were utilized in three distinct but complementary ways: first, to constrain (optimized and non-optimized) model parameters; second, to define target values for optimized model features; and third, to define further quantitative tests that we used to validate the behavior of the optimized models in diverse paradigms. Optimization and validation targets were kept fixed during model development, but our assumptions about model parameters were revised several times based on the results of model optimization and validation.

There was substantial diversity in the recordings available for the extraction of the feature values used as the targets for optimization and somatic feature validation. Since the target values correspond to the mean of the experimental features (with the error scale defined by the standard deviation), including qualitatively different recordings (with a possibly multimodal distribution of some features) could result in large standard deviations and mean feature values that are far from the feature values that characterize any of the recorded neurons. To avoid this situation, we excluded some atypical recordings, resulting in experimental data that were qualitatively similar but still sufficiently diverse at the quantitative level.

The diversity of the recordings (that remained even after removing outliers) also caused some technical issues that we needed to handle. In particular, we encountered cases where a given current injection evoked a spike (or several spikes) in some recordings but no spike in other recordings (from the same or a different neuron). In these cases, the mean values of action potential features across all the recordings at this current amplitude could not be defined in a meaningful way, and we evaluated models only on the features that were valid for all experimental recordings. On the other hand, when all the experimental traces for a certain stimulus contained action potentials but the model did not fire and thus errors in spike features could not be computed (and did not contribute to the total error), we added a large penalty (+100) to the final error score to make sure that models could not achieve a better score simply by not firing action potentials.

To fit the parameters of the models, features from the Electrophys Feature Extraction Library (eFEL) were used instead of using the voltage traces directly, which makes parameter fitting more reliable and robust [90] [91] [1] [2]. eFEL is a validated and widely used library that contains a large variety of features, which allowed us to cover many aspects of the voltage response rather than focusing on a few hand-picked features that could easily lead to overfitting. However, even with this wide range of features, sometimes we still encountered model behaviors that seemed acceptable based on the feature values and resulted in a small overall error score, but a trained human observer could immediately see that something was amiss. In one case, the size and shape of the afterdepolarization in the tuned models were clearly different from the data, but this was not properly captured by the features calculated by eFEL. In another case, the best-fitting models matched the experimental spike pattern by producing a visible calcium spike in the somatic trace, although there was no sign of a slow calcium spike in the experimental recordings, and it is known that CA1 PCs do not tend to produce Ca spikes in response to weak somatic current injections [92]. In these cases, after carefully evaluating the causes of the strange behavior, the model’s boundaries needed to be adjusted in a way that might not lead to an improvement in the error scores. In some cases, we also needed to adjust boundaries when the optimization algorithm consistently found solutions that were very close to one edge of the allowed parameter range, which could indicate that our initial guess regarding the plausible range of that parameter was incorrect, and extending the range could lead to better solutions.

Last but not least, optimizing the model for somatic features does not constrain parameters that have more impact on the dendritic behavior but have little effect on what is happening at the soma. To obtain realistic results for dendritic behavior, the properties of the channels that strongly affect dendritic excitability (mainly the SK-channels and R-type calcium channels) or the boundaries corresponding to their densities (which were optimized parameters) had to be adjusted, and the parameter fitting based on somatic behavior was repeated.

In conclusion, even with our current workflow relying on automated tools, building good models still needs a lot of human input, and several rounds of optimization and validation may be necessary, which can be rather time-consuming both in terms of computations and human effort. One solution could be an extended multi-objective approach in which somatic features and dendritic behavior can be fitted at once, but this would heavily increase the computational demands, and new tools should be developed, or the current tools should be extended to support this scenario.

### The performance of models on the tests of HippoUnit: challenges and limitations

We used five previously created tests implemented by the HippoUnit package [16] to evaluate models under several different conditions. The final models that we created achieved good scores on four of the five tests, while most of the models failed the Depolarization Block test. This test is designed to capture the experimental observation that sustained strong depolarizing input drives CA1 pyramidal cells (and many other types of neuron) into a state where, following a few initial spikes of decreasing amplitude, they stay depolarized but are unable to fire full-amplitude somatic action potentials. Several in vitro studies, including the one from which the target data for HippoUnit’s Depolarization Block Test originated [20], reported an essentially flat membrane potential during depolarization block, while others showed smaller somatic spikelets that likely reflected axonal or dendritic spikes that failed to induce full amplitude somatic action potentials due to the sustained inactivation of somatic sodium channels [61]. Our models consistently showed the latter behavior, which was inconsistent with the results of Bianchi et al., and caused most of the models to fail this test even though their behavior was consistent with the results of other experimental work. The few models that passed the test had qualitatively similar behavior but slightly smaller spikelets whose amplitude was below the detection threshold of the test. The cause of these qualitative differences between different experimental studies is currently unclear, but differences between in vitro and in vivo conditions, and also in recording conditions (temperature, electrodes, solutions) between different in vitro studies, are likely candidates.

We found that the spikelets during depolarization block in the model are caused by full amplitude action potentials in the axon, and the axon (and especially its initial segment) has been identified as a possible source of somatic spikelets in other studies as well [61]. Therefore, we expect that the (morphological and biophysical) properties of the axon will have a large impact on the generation of spikelets during depolarization block, and a more realistic model of the axon could be used to investigate this matter further.

The resting membrane potential of neurons can vary significantly even within a given cell type, and it can also change as a result of synaptic inputs, neuronal activity, or experimental manipulations (including the effects of recording electrodes). The recordings that we used as the target for the Somatic Features Test standardized the baseline voltage by injecting a constant current to achieve a membrane potential around -65 mV before the injection of current steps. This experimental manipulation was not replicated in our simulations; instead, a resting potential around -65 mV was among the target features for parameter optimization, and this target was successfully achieved in all of our optimized models.

A potentially related issue regarding the Oblique Integration Test was that, in many cases, combinations of synaptic inputs to oblique dendrites that triggered supralinear dendritic integration also evoked somatic action potentials, which had a dominant effect on the amplitude of the somatic voltage response. The target data from Losonczy et al. did not seem to contain somatic APs, although it was not clear whether this was due to any filtering or manipulations (such as current injections). Therefore, we injected a small negative constant current into the somata of our models while running the Oblique Integration Test, which significantly reduced the occurrence of somatic spikes and increased the number of dendritic segments from which meaningful results could be obtained.

Validating our optimized models also revealed one shortcoming of the Oblique Integration Test of HippoUnit. We observed that some models that achieved good scores on most features of this test and whose behavior also appeared acceptable upon visual inspection of the voltage traces, nevertheless received a high error score on the “time to peak at threshold” feature. We found out that this was due to the relevant voltage signal having two components: the fast sodium spike component and the slower NMDA component. HippoUnit currently measures the latency of the highest peak, which may correspond to either the fast or the slow component, depending on the overall shape of the voltage response of the model, while the data from the study of Losonczy & Magee 2006. [17] always refer to the latency of the fast component. This results in an artificially inflated error for models where the overall highest voltage is achieved during the NMDA component of the response. This problem could be solved by modifying the Oblique Integration Test of HippoUnit to separate the two components of the signal and always measure the timing of the first component.

One important general conclusion of our study is that building high-quality general-purpose models requires the evaluation of model behavior in diverse settings. HippoUnit currently consists of five tests that validate different aspects of CA1 PC physiology, and such diverse testing was critical for sufficiently constraining the unknown parameters of the model. For example, we found that optimized CA1 PC models with NMDA receptor models from previous studies could achieve good scores on the Somatic Features Test, the PSP Attenuation Test, and the Oblique Integration Test if the density of dendritic Na channels was assumed to be relatively high; however, these models performed badly on the Backpropagating Action Potential Test. On the other hand, by modifying the voltage dependence of the NMDA receptor model and using a lower density of dendritic Na channels, we could build models that performed well simultaneously on all of these tests. Finally, although we believe that using the current tests of HippoUnit one can constrain most parameters well, and build models that generalize to novel situations, extending HippoUnit with more tests could lead to further improvements in model quality and allow the verification of model behavior in other relevant scenarios.

### Further modelling opportunities

#### Morphology

Nowadays, many reconstructed rat CA1 pyramidal cell morphologies are available, and several recent modelling studies used more than one morphology to strengthen their conclusions. For this study, we used only one morphological reconstruction from Megías et al 2001 [93] because of the high computational demands of parameter optimization and validation, especially for the more detailed versions of the models. However, it would be interesting to see how using different morphologies influences the results. We also note that systematic errors in the morphological reconstructions are expected to have a major impact on the accuracy of the modeling results, so any extension of this work should be based on morphologies that are similarly accurate, especially regarding dendritic diameters.

#### Ion channels

Ionic currents in our model are described by a conductance-based formalism or, in the case of Ca currents, by a permeability-based approach captured by the Goldman-Hodgkin-Katz equations. In both cases, the open probability of the channels is described by the formalism introduced by Hodgkin and Huxley [94], which essentially assumes that the opening and closing of channels is due to the operation of several, independently operating gates, and the open probabilities of individual gates (the gating variables) follow first-order kinetics with transition rates that may depend on the membrane potential or other variables. Among several possible parameterizations of the gating equations, one that is closely tied to the results of typical voltage-clamp experiments was suggested by Borg-Graham [95]. We used this formalism to describe the gating of most channels in our model, and set the parameters to capture the experimentally measured voltage dependence of the steady state value and time constant of channel activation and inactivation.

However, recent experimental data indicate that these types of channel models may not fully capture the behavior of certain ion channels; for instance, channel activation and inactivation may not be independent processes [96] [97] [98]. A more general formalism for describing ion channels is based on Markov chains, which explicitly model the transitions between their states. There are several ion channels in CA1 PCs for which implementations as Markov-type models are available [99] [100] [101]. However, the parameters of Markov-type models are not directly related to electrophysiological measurements, and may not even be sufficiently constrained by the data that are available for most channels. Markov-type channel models also tend to require more computational resources than Hodgkin-Huxley-type models. For these reasons, we used Hodgkin-Huxley-type models of all voltage-gated channels in our model. On the other hand, it would be interesting to see how the model’s behavior changes if certain ion channels are changed to potentially more accurate Markov-type channel models.

Knowing the exact molecular composition of the channels responsible for ionic currents can potentially lead to better neuronal models. However, building a cellular model based on molecularly identified ion channels would require not only knowing the identities and total amounts of all the relevant ion channel proteins, but also their spatial distribution within the neuron, the exact ways they combine with other subunits (including modulatory ones), and the functional properties of the resulting channels that could be cell-type- and even state-dependent. All the required information is typically not yet available, and building models based on partial information can be counterproductive. For instance, we attempted to further improve our model by separately modeling the three molecular subtypes of T-type Ca channels that were detected in CA1 PCs. This approach seemed promising since we found quantitative data on the distribution of the three relevant channel proteins (Cav3.1, Cav3.2, and Cav3.3) [102], and another study described the voltage-dependent gating of Ca channels containing these subunits (in … cells) [103]. However, these molecularly identified channels generated currents whose properties were different from those of T-type currents previously measured in CA1 PCs, and including these channel models in our cell model (and optimizing its parameters) led to unrealistic subthreshold physiological behavior. The discrepancies could be due to the cell-type-specific molecular composition and modulation of the channels, but more data (preferably from molecularly characterized channels in CA1 PCs) will be required to resolve the issue.

On the other hand, we indirectly used molecular information several times when we constructed our model. Modelling sodium channels, knowing that the dendritic, somatic, and axonal sodium channels have different kinetics (due to the distinct spatial distribution of Nav1.6 and Nav1.2 channel proteins in the cell) [25], improved the behavior of the model. Similarly, knowing that D-type potassium currents are generated by a heteromer of potassium channels, which, as homomers, mediate delayed-rectifier potassium currents [26] implied that a D-type potassium current may exist on the axon as well, which also improved our model.

There are also ion channels that were excluded from this model, but, following the ever-growing literature, it could be an interesting experiment to determine how including these channels would alter the model’s behavior. One of the candidates would be the inward-rectifier potassium channels that are also present in the dendrites and dendritic spines and influence the excitability of the cells [104] and can play a role in synaptic plasticity mechanisms. [105] Modelling chloride channels are also a possible direction for adding further ion channel models since they play an important role in maintaining the balance between excitation and inhibition, shaping synaptic responses [106], and protecting against pathological conditions like ischemia and epilepsy. [107]

## Methods

### Data

To construct realistic models of hippocampal CA1 pyramidal neurons and validate their somatic electrophysiological behavior, we used somatic whole-cell patch-clamp recordings from rat CA1 PCs. The data were provided by Judit Makara and were previously described in Sáray et al. (2021). [16] The recordings were obtained in current-clamp mode and contain the somatic membrane potential responses of 6 CA1 PCs (with 3 measurements from each cell) to somatic step current injections of 300 ms duration and ten different amplitudes (-0.25, -0.2, -0.15, -0.1, -0.05, 0.05, 0.1, 0.15, 0.2, and 0.25 nA). One of the 6 cells displayed a very different voltage response from the rest, so we excluded it from the target data. A typical experimental voltage response can be seen in Figure 1C. From the raw voltage traces, we extracted electrophysiological features using BluePyEfe (https://github.com/BlueBrain/BluePyEfe) and the eFEL feature extraction library (https://github.com/BlueBrain/eFEL) [108].

For those features that average over all the action potentials we always excluded the first action potential because of a systematic error in the detection of action potential initiation near the beginning of the current step. Sag features were evaluated only for negative amplitudes, voltage deflection features were only considered at non-spiking amplitudes, while spiking features were evaluated for current steps where spikes could be observed in the experimental data.

For parameter fitting, we selected a set of eFEL features that best describe the somatic behavior of the cell with minimal redundancy. The selected features are listed in Table 2. For validation of somatic behavior, a wider range of features was used, as described in Saray et al. 2021 [16].

### Model building

#### Morphology

We used morphology pc1a from Megías et al. 2001. [93], which had reconstructed soma and dendrites in detail, and we also added a few axonal compartments from the model of Bianchi et al. 2012 [20]. The reconstructed morphology can be seen in Figure 1B. Megías et al. also give detailed information about the number of dendritic spines of each dendritic region, from which we calculated the spine density (number /µm length) for the basal dendrites (2.85), trunk (4.52), oblique (3.52) and tuft dendrites (0.56), and the distance between spines on a dendritic section was 1/density in the corresponding region. The dimensions of all dendritic spines were modelled according to Harnett et al. 2012 [109] (spine neck length: 1.58 µm, spine neck diameter: 0.077 µm, spine head length and diameter: 0.5 µm).

With this method, we can explicitly model any number of spines at any location, taking the spine density of the region into account, including modelling all the possible spines on the morphology, resulting in a total of 29602 dendritic spines. However, modelling all dendritic spines greatly increases the computational cost of the model, and therefore, in most of our simulations, we used a method that takes spines into account in a computationally more efficient way.

There are several different approaches to take the dendritic spines into account without explicitly simulating them, but the most principled and computationally efficient way is the correction of the appropriate biophysical parameters with the surface factor (F-factor). The F-factor is defined as the relative increase in the membrane area of a dendritic section due to the presence of spines. The spines consist of a spine neck, which is modeled as a cylinder, and a spine head, which can be modeled as a sphere or as a cylinder. We note that if both the length and the diameter of the cylinder are equal to the diameter of the sphere, they have equal surface areas (excluding the bases of the cylinder). Therefore, for convenience, we decided to model the spine head as a cylinder. From this, the surface of the spine can be obtained as follows:

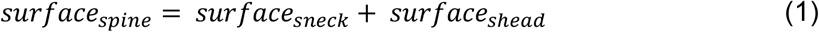

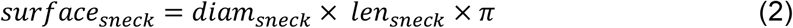

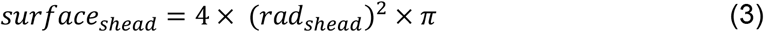

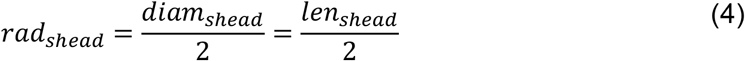

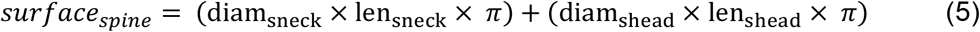

The surface area of a dendrite of length *L*_*dend*_ and *diam*_*dend*_ is

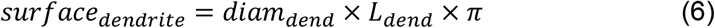

and the total surface of the spines on that dendrite is

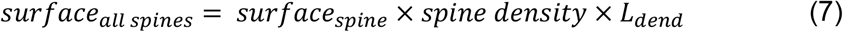

From this the F-factor for that dendritic segment is:

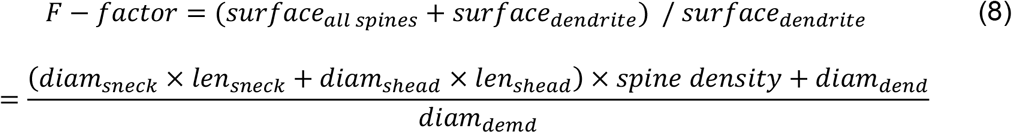

Then, instead of modeling each spine explicitly, their effect on electrical signaling in the dendrites can be taken into account by multiplying the leak conductance and the membrane capacitance by the corresponding F-factor. A similar approach can be applied to active conductances that are present on the spines [83]. The F-factor was calculated independently for every section of the dendritic tree. For every dendritic section where we did not explicitly model them, we take the spines into account by the F-factor. For those dendrites that wear explicitly modeled spines (any number), the F-factor was calculated with the exclusion of the membrane area of the modeled spines.

#### Passive parameters

The specific membrane capacitance was set to 1 µF/cm^2^, which corresponds to the standard experimentally measured value for the neuronal membrane and was later multiplied by the F-factor on the affected dendritic sections. The specific axial resistance (Ra) of the model was set to 50 Ohm×cm, which is lower than the values typically used in previous models of CA1 PCs. This lower Ra is in alignment with recent model-based estimates of passive parameters of CA1 PCs [60] and was essential for the model to capture a wide variety of experimental data on signal propagation and integration.

#### Channel models

The model contains 13 active voltage- and/or calcium-gated ion channels, and occasionally 2 or 3 versions of one type of ion channel when the kinetics of the ion channels are dependent on their distances from the soma or on the dendritic region in which they are located. Our channel models are diverse in implementation and in their sources, but every one of them was carefully evaluated in the light of the newest data that can be found in the literature about their kinetics and were modified accordingly when necessary.

The current that flows through an ion channel at a given membrane potential can be given as:

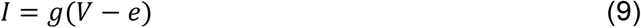

where *V* is the membrane potential, e is the equilibrium potential or reversal potential of the given ion, and g is the conductance of the ion channel. The voltage and/or calcium-dependent g is the product of the theoretical maximal conductance of the ion channel and gating variables independent from each other, corresponding to the number of activation and inactivation gates that the given ion channel possesses. A gating variable *X*, can be described with the following equations based on the Hodgkin-Huxley model:

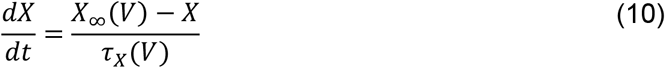

where *X*_∞_is the steady-state value of *X, τ*_*X*_ is the time constant, and both are dependent on the voltage (or the calcium concentration in some cases).

*X*_∞_ and *τ*_*X*_ can both be described by several different formalisms. In the more traditional Hodgkin-Huxley equations, they are defined in terms of the forward (α) and backward (β) rates of channel opening and closing. In this formalism

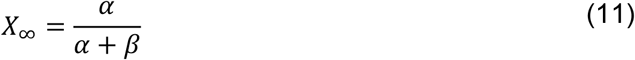

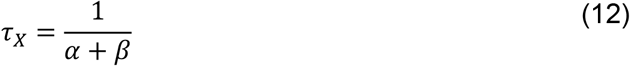

Although for some ion channels we used models that employ this kind of equation, experimental data regarding ion-channel kinetics rarely provide these rates directly. Instead, voltage-clamp experiments typically measure the voltage- and /or calcium concentration-dependent steady state activation and inactivation as well as the associated time constants. For easier parameterization and to adjust our channels more directly to the experimental data, we described most of our channels using the formalism introduced by Borg-Graham [110], where:

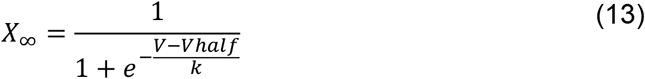

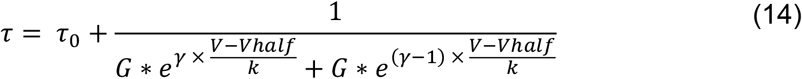

where *Vhalf* is the voltage where *X* is half-activated at steady state, 1 /(4k) is the slope of the steady-state activation curve at *Vhalf, G* is a rate coefficient and *γ* is an asymmetry parameter (where 0 =< *γ* <= 1) for the voltage-dependent part of *τ*, and *τ*_0_ is the voltage-independent part of *τ*. From these equations:

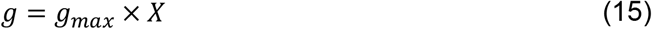

if the channel has only a single activation gate. Additional gates (for activation and inactivation) can be described as multipliers of the right-hand side of this equation.

The models of calcium channels use the Goldman-Hodgkin-Katz equation to describe the current, which gives a better fit for currents mediated by calcium ions than the traditional conductance-based Hodgkin-Huxley formalism. In this description, instead of the conductance, the theoretical maximal permeability is used, and the activation and inactivation curves are multiplied by the GHK equation instead of the driving force. The GHK equation looks like the following:

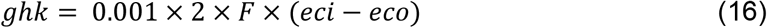

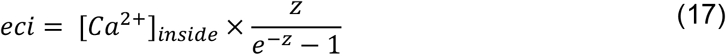

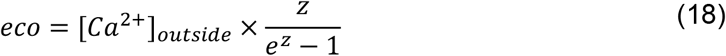

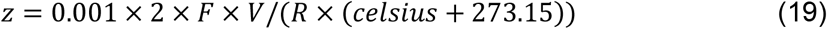

where *F* is the Faraday constant, and *R* is the gas constant, and *celsius* is the temperature of the model is in degrees Celsius. From this

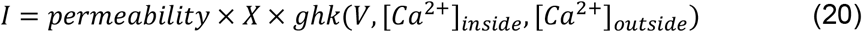

The value of [*Ca*^2+^]_*outside*_ is the default value for the extracellular calcium concentration of the Neuron simulator (2 mM) while the [*Ca*^2+^]_*inside*_ is determined by the model of the calcium dynamics (see later). Since we tried to find and, in many cases, create ion-channel models that best describe the latest experimental results, differences from these equations can occur. In those cases, we state the equations and parameters in the description of the ion channel in question.

In the following description of the channel models, the voltage dimension values are expressed as millivolts (mV) while the time dimension values are in milliseconds (ms).

##### Hyperpolarization-activated mixed cation (Ih) current

The model of the h-current was based on the observations and measurements of Zemankovics et al. [21]. Since the earlier data of Magee [22] indicated that the proximal and distal channels differ in the voltage-dependence of their activation kinetics, two different channel models were incorporated into the model and distributed throughout the dendritic arbor in a non-uniform fashion [22].

For the proximal H-current:

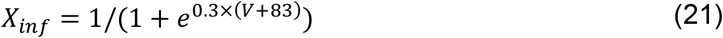

For the distal H-current:

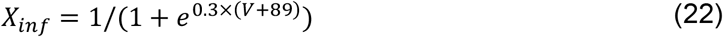

and for both channels the τ was:

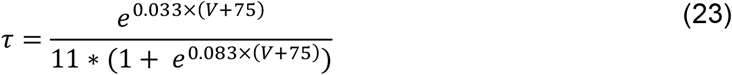

To determine the exact amount of Ih in the model, a ratio parameter was fitted during the optimization process that served as a multiplier in each dendritic segment. With this ratio parameter, the equation for the non-uniform distribution from 0 to 100 µm distance from the soma is:

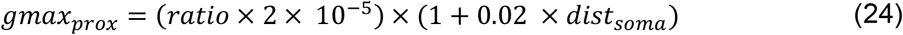

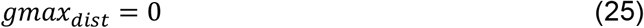

From 100 µm to 400 µm:

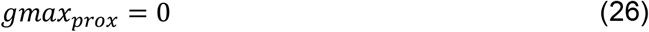

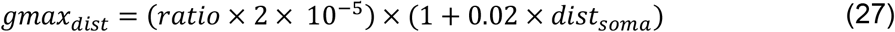

and further from 400 µm:

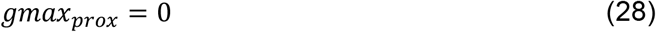

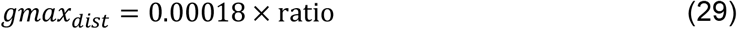

Sodium channel

The model has 3 types of sodium channels that have modified kinetics for the soma, axon, and dendrites.

Sodium current was given as the following:

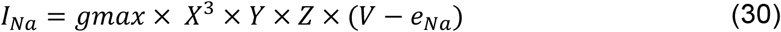

*gmax* is the maximal conductance of the channel, *X* is the gating variable for activation, *Y* is the gating variable for fast inactivation, *Z* is the gating variable for slow inactivation, *V* is the voltage, and e_Na_ is the reversal potential of the sodium channel. *X* and *Y* were described in the Borg-Graham formalism stated above (Eq. 13-14). The steady-state half-activation and half-inactivation were fitted parameters. To ensure that the action potentials are initiated in the axon, both the half-activation and half-inactivation were more hyperpolarized by 3 mV than those of the sodium channels in the soma and dendrites. [24]

The characteristics of the slow inactivation gate are based on data from Colbert et al. 1997. [111] and Jung et al. 1997. [112]. In particular,

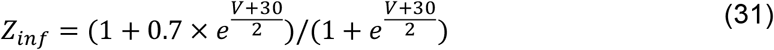

for the somatic sodium channels, and

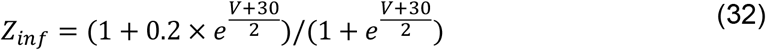

for the dendritic sodium channels. The expression for *τ* is the same for both somatic and dendritic channels:

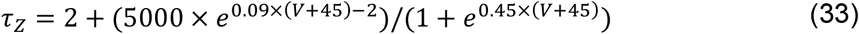

The time constants were provided for CA1 pyramidal neurons by Martina & Jonas 1997. [113]

The somatic density of sodium channels was a fitted parameter. The axonal sodium channel density was 40 times the somatic density, while on the oblique and basal dendrites it was 2/3 of the somatic density, and on the tuft dendrites it was 1/3 of the somatic density, based on the data of Lőrincz and Nusser 2010 [25]. On the apical trunk, the somatic density decreased with distance according to the equation

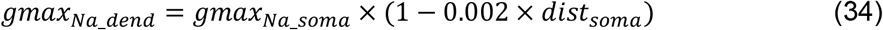

##### Potassium channels

###### Delayed rectifier potassium channel

Delayed-rectifier potassium channels contribute significantly to action potential repolarization and therefore have a major impact on the shape and height of the APs, along with the sodium channels. [114]. They were modeled according to the Borg-Graham formalism described above (Eq. 13-14). The parameters of the equations can be found in Table 3. Although this channel can be inactivated according to experimental data, this process is so slow that it is not expected to be significant under physiological conditions and was therefore not modeled.

**Table 3:**
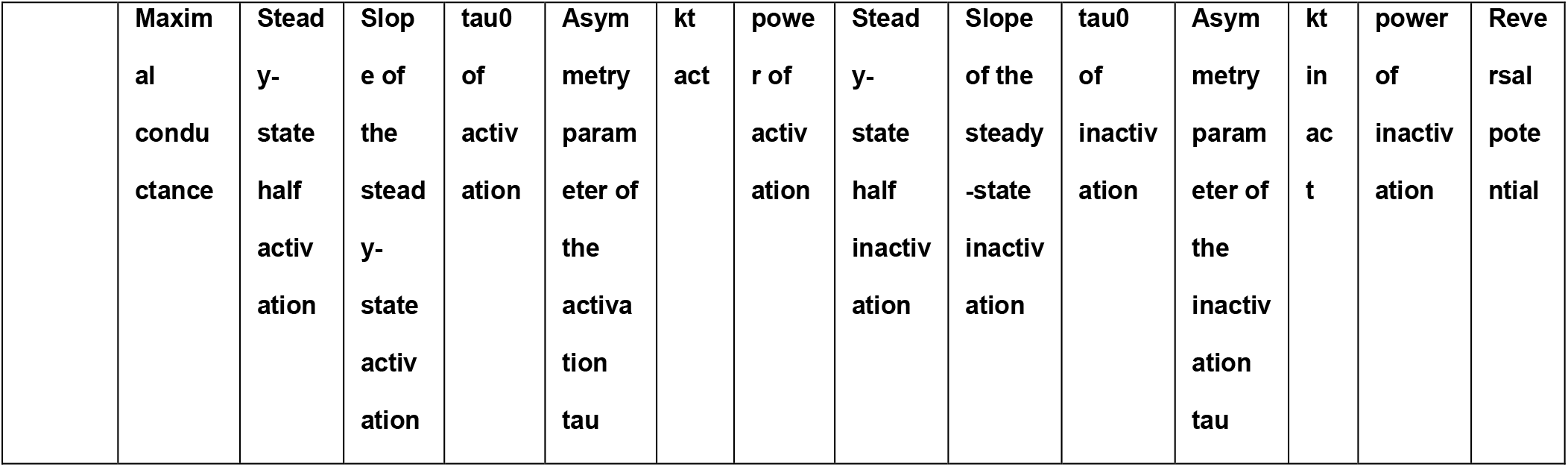
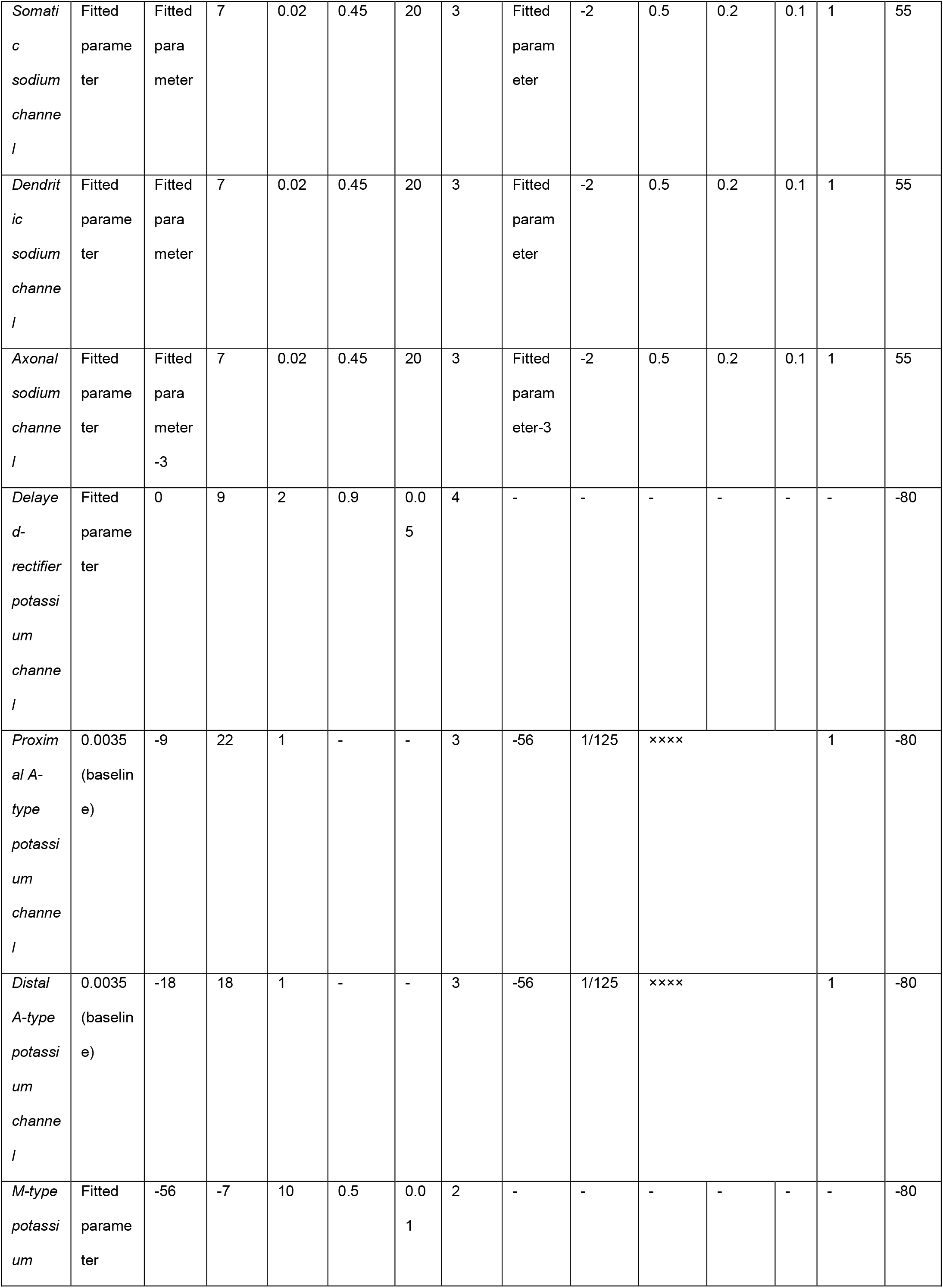
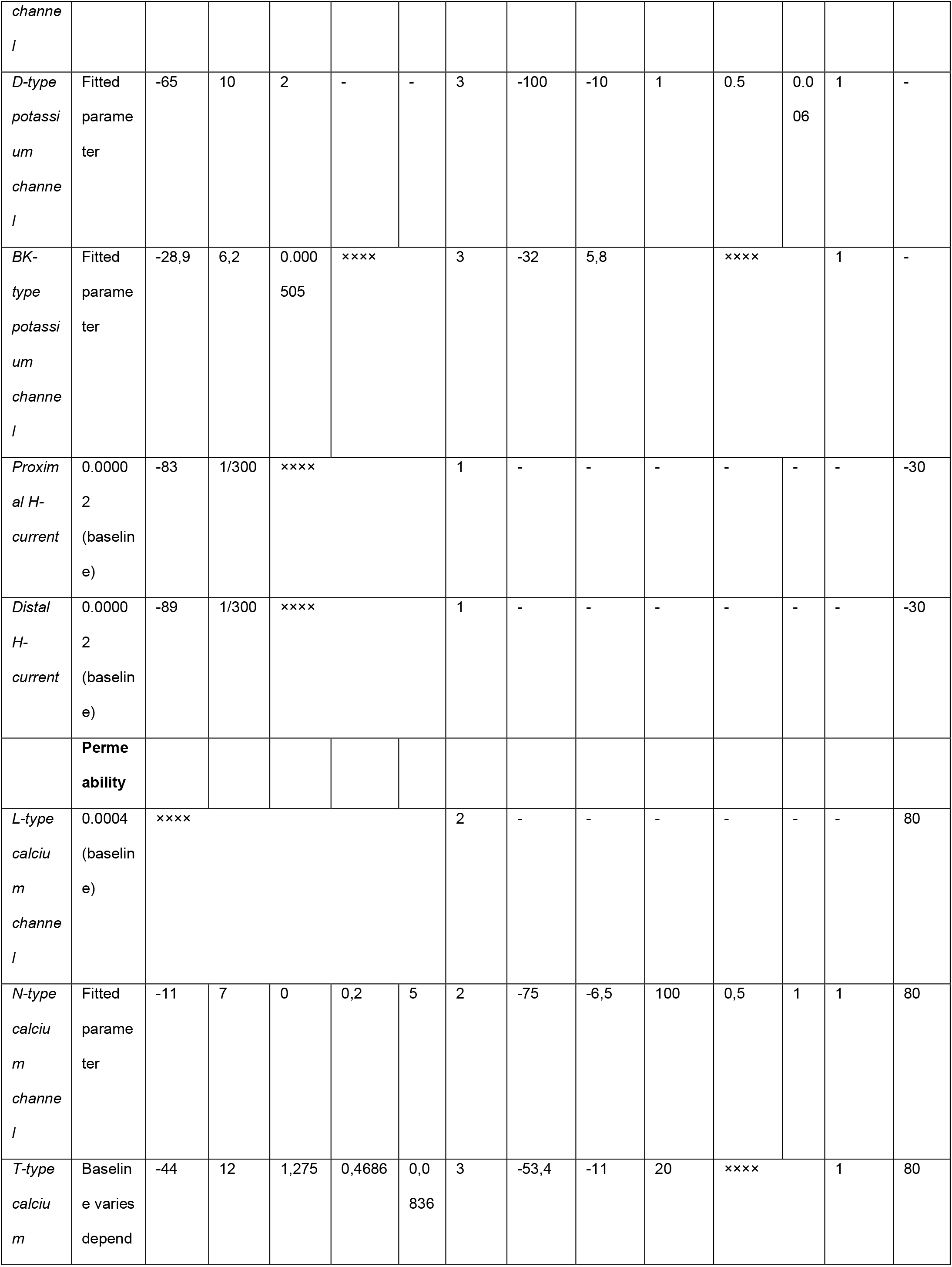
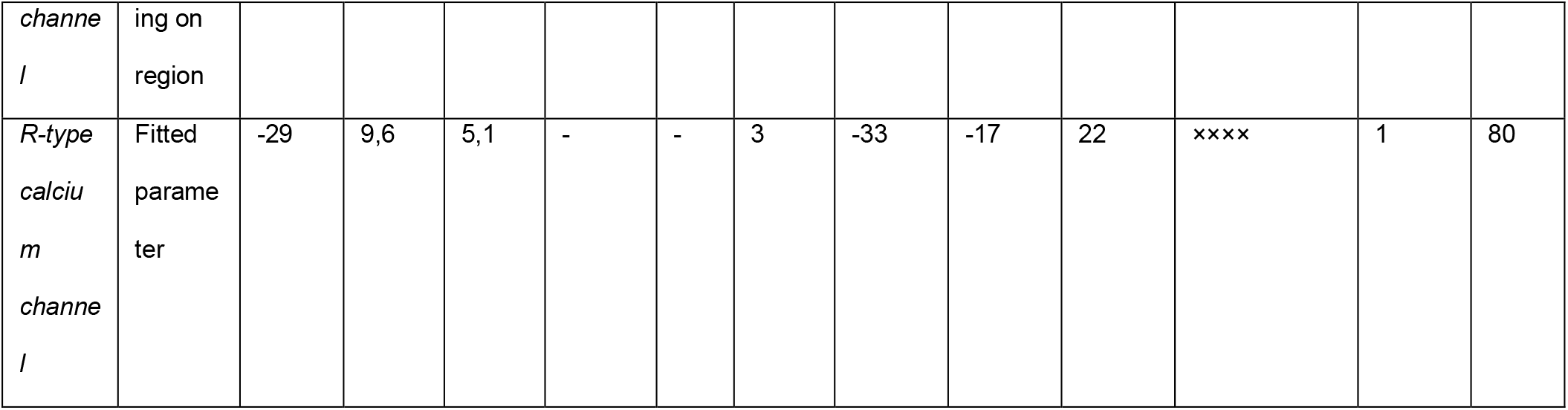
Parameters of the ion channels present in the model. Cells containing ×××× have their equations in different formalisms; for these equations and parameter values, see the corresponding parts of Methods.

###### M-type potassium channel

The M-type potassium channel has roles in generating the somatic medium afterhyperpolarization, preventing bursting, and generating subthreshold resonance in the theta frequency range. It also plays a major role in determining the action potential threshold and suppressing the afterdepolarization.

We used the channel model from Hönigsperger et al. 2015. [32] The channel model uses the standard Borg-Graham formalism (Eq. 13-14). The parameters of the equations can be found in Table 3. We used the same maximal conductance on the soma and the dendrites, which was a fitted parameter, while on the axon, the maximal conductance was 4 times higher than the somatodendritic value. [74]

###### A-type potassium channels

A-type potassium channels play key roles in the regulation of dendritic excitability [34] [115], but they also contribute to action potential repolarization [116] and influence subthreshold neuronal behavior [115]. Our model of the transient A-type potassium channel was based on the observations of Hoffmann et al. 1997. [34] They found that the voltage dependence of its activation was different in somatic and dendritic regions of the cell. We implemented two types of this channel, one for the soma and the proximal dendritic regions, and one for the distal dendritic regions. The voltage-dependent activation and inactivation of K(A) in the model were also based on data from Hoffmann et al. (1997), but we allowed some flexibility in a few specific parameters to make sure that this channel could fulfil all the diverse roles that are indicated above. In particular, the voltage dependence of the steady-state activation and inactivation of the channel was described with the Borg-Graham formalism (Eq. 13-14), and its baseline parameters can be found in Table 3. However, we allowed the slope of voltage-dependent K(A) activation to change during parameter optimization, and we also introduced an additional free parameter that shifted the half-activation and half-inactivation voltage by equal amounts.

The *τ* of the channel activation was set to a voltage-independent constant of 1 ms [34], while according to the same article, the tau of inactivation above -30 mV

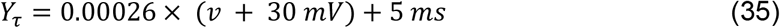

and below -30 mV

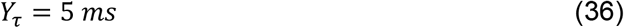

Hoffmann et al. also stated that the maximal conductance of the A-type potassium channels is linearly increasing with the distance from the soma up until 400 µm. Since we have no data further from that point, we set the maximal conductance constant in the distal dendrites. The model follows this inhomogeneous distribution by the following:

On the soma, we have

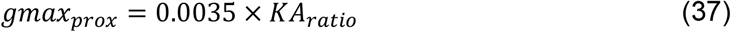

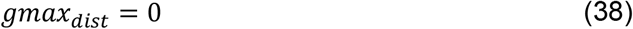

from the soma to 100 µm distance

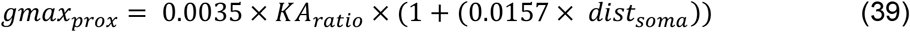

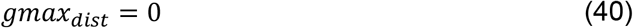

from 100 µm to 400 µm

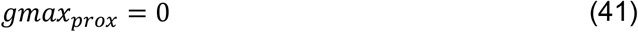

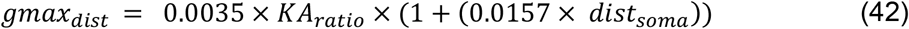

and above 400 µm

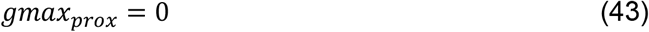

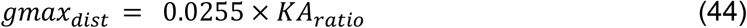

The value 0.0035 from the maximal conductance comes from Kerti et al. 2011. [117], and *KA*_*ratio*_ is a fitted parameter.

###### D-type potassium channel

The D-type potassium channel is present on the soma and the axon, and its kinetics were carefully examined and adjusted to the literature to get the delay at the beginning of the current step to the first action potential, which can be observed in the experimental data. The channel model is defined by the Borg-Graham formalism (Eq. 13-14), and its parameters can be found in Table 3.

The channel model originates from Golomb et al. 2007. [118] [119] [30] [31] The steady-state half-activation and half-inactivation were changed from -60 to -65 and from -90 to -100, respectively.

The parameter gkd, which is the maximal conductance of the channel, is a fitted parameter. Big-conductance (BK) calcium- and voltage-dependent potassium channel

Big-conductance calcium and voltage-dependent potassium channels are located on the soma, co-localized with N-type calcium channels [44], so in the model, it takes its calcium input solely from what comes in through the N-type calcium channels. It has a voltage-dependent activation and inactivation and a calcium-dependent component that are modelled by three separate gates. The voltage-dependent gates were described with the Borg-Graham formalism (Eq. 13-14, parameters in Table 3), and the calcium-dependent gate looked like the following:

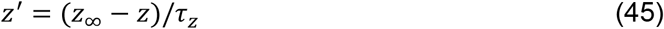

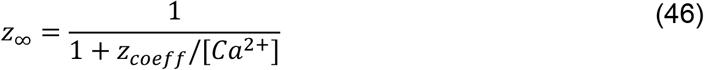

where *z*_*coeff*_ = 0.001 mM, *τ*_*z*_ = 1 ms. From this the potassium current coming from this channel is:

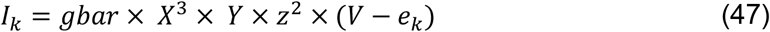

###### Small-conductance (SK) calcium-dependent potassium channel

Small-conductance calcium-dependent potassium channels are present on the soma and dendrites. SK channels are also present in the spine heads [54], and there is an approximate number of how many channels are in one spine (2-3 channels/spine). The number of SK channels in dendritic membrane patches and how this number is changing in the different dendritic regions was also available. It was a linear change, so we implemented that in the model, and we left the somatic value (from where it will linearly increase) as a free parameter between boundaries close to the experimental data. [54] Since this is a calcium-activated potassium channel and has no voltage dependence, the equations are different from the other potassium channels that are voltage dependent. The equations came from the channel model of Hirschberg et al. [55]. This channel model also had a built-in temperature dependence that we decided to use because the experimental data that it was fitted to was measured at room temperature (23 C) while our experimental data was measured at body temperature (33 C).

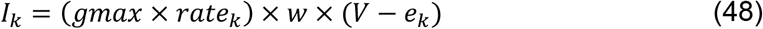

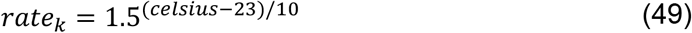

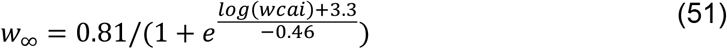

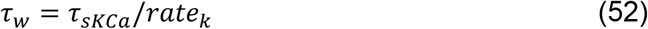

where *τ*_*sKCa*_ = 2.365 ms and *wcai* is the calcium concentration.

The *gmax*_*sKCa*_*soma*_ was a fitted parameter. We fitted an equation to get the maximal conductance ratios equal to the SK channel densities in different dendritic regions stated in Ballesteros et al. 2012. [54] From this

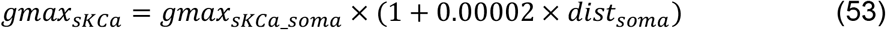

###### Slow Afterhyperpolarization potassium (K-AHP) current

The potassium current responsible for the slow afterhyperpolarization that occurs in the CA1 pyramidal cells was initially assumed to be generated by the SK-channels, but this later proved to be false. There were also reports about intermediate-conductance calcium-dependent potassium channels that are present in other cell types, may also exist in CA1 pyramidal cells, and may be responsible for the slow afterhyperpolarization. [56] Since there was a debate about these findings and there is no available data about the kinetics of this channel, we kept a phenomenological calcium-dependent potassium channel model that produces the desired behavior at the soma. It was described with first-order kinetics modeled using transition rates, similar to the original Hodgkin-Huxley equations (Eq. 11-12), where α = 0.2× calcium concentration if the concentration was less than 50 mM, and α = 10 if the concentration was above 50 mM, and β = 1. The maximal conductance (gmax) was a fitted parameter. Since the channel that is responsible for slow afterhyperpolarization is co-localized with the L-type calcium channels [44], our channel model gets the calcium ions solely from the L-type calcium channels. The kinetics were based on the observations of Traub et al. 1991. [57]

##### Calcium channels

###### L-type calcium channel

High-voltage-activated L-type calcium channels play a major role in shaping the currents in the dendrites and spines of the CA1 pyramidal cell. Although both calcium and voltage-gated inactivation exist in these channels, they were neglected because of their slow time course. The steady-state activation is based on Bell et al. 2001 [46], and the current is described by the GHK equation (Eq. 16-20).

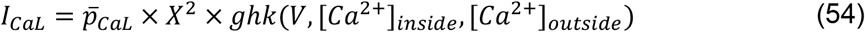

The rates are the following:

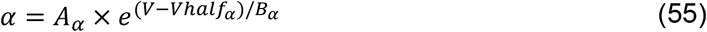

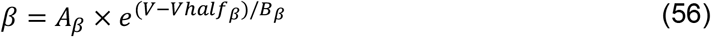

where *A*_*α*_ = 0.3, *B*_*α*_ = 17 and *Vhalf*_*α*_ = 6.5, while *A*_*β*_ = 0.3, *B*_*β*_ = - 17 and *Vhalf*_*β*_ = 6.5 The permeability of the channel was a fitted parameter. According to Magee et al. 1998. [45] and Marion and Tavalin 1997. [44] L-type calcium current is present perisomatically, then decreases further from the soma; therefore, we applied an inhomogeneous distribution of this ion channel with the equation of

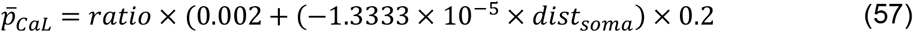

until 150 µm distance and from then

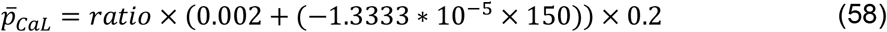

The fitted parameter was the ratio.

###### N-type calcium channel

The model contains N-type calcium channels exclusively on the soma. Channel gating is described by the classical Borg-Graham formalism, and the current is calculated via the GHK equation (Eq. 16-20).

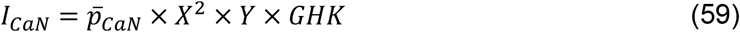

The parameters for the *X* and *Y* equations can be found in Table 3. The kinetics were based on McNaughton and Randall 1997. [42]

###### T-type calcium channels

We decided to implement one T-type calcium channel that describes the current measured in a CA1 pyramidal cell [47]. The steady-state activation and inactivation curves and the time constant of the activation were described by the Borg-Graham formalism (Eq. 13-14), whose parameters can be found in Table 3, while the time constant of the inactivation was described as follows:

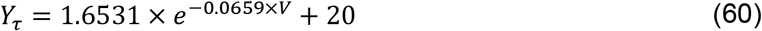

The number of different T-type calcium channel isoforms was explicitly measured by McKay et al. [48] in every region of the dendritic arbor and the soma. Since the model has only one type of T-type calcium channel, we summed up the total numbers of each isoform present in the given region. We could not find a simple fitting equation to describe the gradient of the current, and therefore, we explicitly used the measured permeability in each region and introduced a single multiplier that scaled all permeability values as a fitted parameter.

The baseline permeability on the oblique dendrites was 7.2608×10^-7^, on the basal dendrites it was 6.9983×10^-7^, on the apical trunk 2.4292×10^-7^, on the tuft dendrites 4.3133×10^-7^, and on the soma, it was 2.125×10^-8^ cm/s.

###### R-type calcium channel

Cav2.3, which is the Ca channel subunit responsible for the R-type current in the CA1 pyramidal cells, is exclusively present on dendritic spines. [49] One spine head can contain 5-15 channels [50]; based on this and the known single channel conductance, we calculated the feasible range of permeability values to set the boundaries for this optimized parameter. The properties of the channel were described by Foehring et al. 2000. [51] and Randall & Tsien 1997 [47]. We used the GHK equation (Eq. 16-20) to describe the channel behavior, and its parameters can be found in Table 3.

In those simulations where dendritic spines were not explicitly modeled, R-type Ca channels were included in models of the corresponding dendritic segments, and their density was set using the surface factor (F-factor) – however, since these channels were assumed to be present only in the spines, the multiplying factor for the maximal permeability was F-factor minus one (rather than F-factor, which would imply an equal density of the channel in the membranes of spines and the dendritic shaft).

###### NMDA receptor

To make the model simultaneously perform well on several HippoUnit tests and qualitatively match the experimental data of Losonczy and Magee 2006. [17] we created a new NMDA receptor model. (See Results for more explanation)

The voltage dependence of the Mg-block was fitted with a sigmoid function described by the Boltzmann equation:

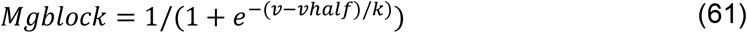

This equation can describe the voltage dependence of many previous NMDA receptor models as well, and its parameters have clear interpretations as the midpoint and (inverse) steepness of the sigmoid. The final parameters are *vhalf* = 6 and *k* = 18.

The equations describing the behavior of the receptor model are:

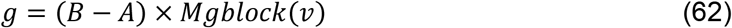

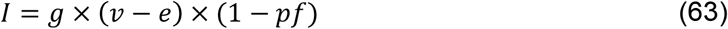

Here, *A* and *B* are state variables that describe the control of presynaptically released glutamate over the opening and closing of the ion channel, *A* corresponds to the rise and *B* to the decay of the current, and *B* − *A* is analogous to *X* in the Hodgkin-Huxley equations, (Eq. 11-12), modified by the synaptic weight of the NMDA channels in the model. Following a presynaptic spike and subsequent transmitter release, A and B change instantaneously according to

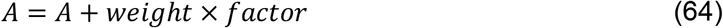

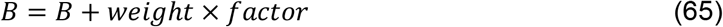

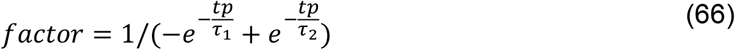

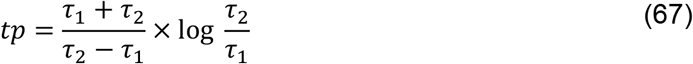

The scaling factor is set such that a single presynaptic release event through a synapse of weight 1 generates a peak conductance of 1.

In the absence of presynaptic activity, A and B decay exponentially according to

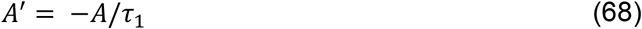

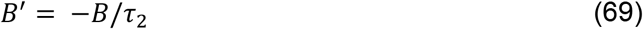

*pf* = 0.03, which is adjusted so that 15% of the whole current through the NMDA channel is mediated by calcium ions at -60 mV, *τ*_1_ is the rise time that equals 3.3 ms and *τ*_2_ is the decay time constant, which equals 102.38 ms. *τ*_1_ and *τ*_2_ are Q10-corrected values from McDermott et al. 2006. [120]

##### Calcium dynamics

The model contains four different types of pools for intracellular calcium ions, each with its own separate dynamics. One general calcium pool is not assigned to any specific calcium channel and is used in the soma and the dendrites, another one is present in the spines, a third calcium pool is for the calcium ions that flow in through the N-type calcium channels and are then used by the BK potassium channels, and a final calcium pool is for the calcium ions that flow in through the L-type calcium channels. The latter is connected to the hypothetical K_AHP current, since the slow afterhyperpolarization is mediated by an only calcium dependent potassium channel that is most likely co-localized with the L-type potassium channels. [44]

The calcium dynamics models calcium accumulation into a volume of area×depth next to the membrane with a decay time constant (tau) to the resting level. For the general calcium dynamics and the dynamics in the spine heads, the equations are the following:

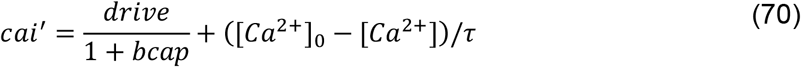

where [*Ca*^2+^]_0_ is the baseline calcium concentration (5×10^-5^ mM), *τ* = 10 ms, and the buffer capacity (bcap) = 17.

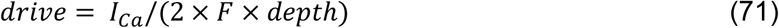

where *F* is the Faraday-constant, and depth is the shell where the calcium ions take effect, most of the time the half of the diameter of the dendrite, soma or spine head, while in the pools assigned to specific calcium channels the *depth* = 1 nm to model the co-localization of the calcium channels and the calcium-dependent potassium channels. If the *drive* <= 0 then *drive* = 0, so it cannot change the direction of the flow of the ions. For the two channel-specific calcium pools *bcap* = 0.

##### Leak

The model also contains a non-specific “leak” current that represents all non-voltage-dependent ionic currents.

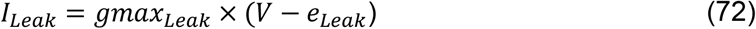

where both the maximal conductance and the reversal potential were fitted parameters of the model.

### Tools

#### NEURON

Simulations were carried out using the NEURON simulation environment. The model was defined in NEURON’s HOC language, and channel models were implemented in the NMODL programming language. [121]

#### Parameter fitting

We used automated parameter fitting methods to determine those model parameters that were not sufficiently constrained by the experimental literature. In a few cases where a published experimental value led to clearly incorrect model behavior, we allowed that parameter to be optimized within realistic boundaries. Table 4 contains the fitted parameters and their boundaries.

**Table 4:** Fitted parameters with their corresponding boundaries.

|  |  |  |
| --- | --- | --- |
|  | Minimum | Maximum |
| Maximal conductance of the leak current | 0.0000005<br>S/cm<sup>2</sup> | 0.00001<br>S/cm<sup>2</sup> |
| Reversal potential of the Leak current | -120 mV | -20 mV |
| The ratio of the inhomogeneous I<sub>h</sub> current from a pre-established baseline | 0.1 | 20 |
| The half steady-state inactivation of the somatic sodium channels | -60 mV | -25 mV |
| The half steady-state activation of the somatic sodium channels | -40 mV | -20 mV |
| Maximal conductance of somatic sodium channels | 0.02 S/cm<sup>2</sup> | 0.5 S/cm<sup>2</sup> |
| Maximal conductance of dendritic delayed-rectifier potassium channels | 0.01 S/cm<sup>2</sup> | 0.2 S/cm<sup>2</sup> |
| Maximal conductance of somatic delayed-rectifier potassium channels | 0.01 S/cm<sup>2</sup> | 0.8 S/cm<sup>2</sup> |
| Maximal conductance of the D-type potassium channels | 0.0001 S/cm<sup>2</sup> | 0.01 S/cm<sup>2</sup> |
| Maximal conductance of axonal D-type potassium channels | 0.000001 S/cm<sup>2</sup> | 0.1 S/cm<sup>2</sup> |
| Maximal conductance of the M-type potassium channels on the soma and dendrites | 0.0000001 | 0.0005 |
| The ratio of the inhomogenous A-type potassium channels from a pre-established baseline | 0.5 | 5.0 |
| A shift in the steady-state activation and inactivation curves of the A-type potassium channels | 0 mV | 50 mV |
| The slope of the A-type potassium channel activation | 5 | 22 |
| Permeability of the N-type calcium channels | 0.002 | 0.01 |
| Maximal conductance of the BK-type potassium channels | 0.001 S/cm<sup>2</sup> | 0.3 S/cm<sup>2</sup> |
| The ratio of the inhomogeneous L-type calcium channels from a pre-established baseline | 0.1 | 5.0 |
| Maximal conductance of the slow afterhyperpolarizing current | 0.00001 S/cm<sup>2</sup> | 0.001 S/cm<sup>2</sup> |
| The ratio of the inhomogenous T-type calcium channels from a pre-established baseline | 0.1 | 5.0 |
| Maximal conductance of the SK-type potassium channels | 0.00001 S/cm<sup>2</sup> | 0.001 S/cm<sup>2</sup> |
| Permeability of the R-type calcium channels | 0.00001 | 0.001 |

The parameter fitting procedure was carried out by the Neuroptimus software [122] which is an interactive Python tool for the automated fitting of unknown parameters between given boundaries. The optimized error function was a sum of standardized feature errors based on the feature values extracted from the voltage traces by the eFEL feature extraction library. For the optimization algorithm, we chose the Covariance Matrix Adaptation Evolution Strategy from the CMAES package as it was found by Mohácsi et al. [14] to be one of the best algorithms to fit the parameters of a morphologically and biophysically detailed neuronal model. We set the parameters of the search algorithm to get the best possible results within an acceptable runtime. The population size was 127, which was determined by the number of cores per node available on the Neuroscience Gateway’s supercomputers [123] that we used for the optimization process. We ran the search for 100 generations, as the results of Mohácsi et al. indicated that this was sufficient for the CMAES algorithm to converge to a solution whose error score was close to the global minimum.

#### Model Testing

To validate the models against experimental data that were not considered during the optimization process, we used the HippoUnit software. [16] HippoUnit is a Python module that automatically performs simulations that mimic experimental protocols on detailed hippocampal cell models. It uses models built in NEURON [121] as a user-defined Python class. We used all the five built-in tests:

The Somatic Features Test uses the eFEL Python package to extract and evaluate the values of both subthreshold and spiking features from the voltage responses of the model to somatic current injections of different depolarizing and hyperpolarizing current amplitudes. Spiking features describe action potential shape (such as AP width, AP rise/fall rate, AP amplitude, etc.) and timing (frequency, inter-spike intervals, time to first/last spike, etc.), while passive features (such as the voltage base or the steady state voltage), and subthreshold features for negative current stimuli (voltage deflection, sag amplitude, etc.) are also examined. Step currents of varying amplitudes are injected into the soma of the model, and the voltage response is recorded. Its outputs are the features extracted from the models and error values resulting from comparisons to the features extracted from the experimental data (see earlier Methods, Data section), an error score that is the mean of the errors of all the features examined by the given test, and figures that illustrate the behavior of the model. This test computes features and errors that are similar to those used during parameter optimization, but it tests more features to determine how the model performs on non-optimized features.

The Backpropagating Action-Potential Test evaluates the strength of action potential backpropagation on the apical trunk at different distances (50, 150, 250, 350 µm) from the soma. The test finds the amplitude of the current injection where the model fires at exactly 15 Hz (or the closest possible rate) and then measures the amplitude of the first and last action potentials in the spike train. [19] The final score here is the average of the Z-scores achieved for the features of first and last action potential amplitudes at different dendritic distances. In the result, it is also stated whether the model is more like a strongly or a weakly propagating cell. [19] We modified the original test from Sáray et al. [16], and instead of evaluating the absolute value of the amplitude of the last action potential, the ratio of the amplitudes of the first and last action potentials was used to address the fact that the shrinkage of the spikes is independent of the absolute value of the spike amplitudes.

The Post-synaptic Potential Attenuation Test tells us about the attenuation of postsynaptic potentials from input sites in different parts of the dendritic tree to the soma. [18] The apical trunk receives excitatory post-synaptic current (EPSC)-shaped current stimuli at locations of different distances from the soma. The somatic and dendritic depolarization values were then averaged in three ranges around distances of 100, 200, and 300 µm from the soma, and the soma/dendrite attenuation was calculated to get the mean and standard deviation of the attenuation features at the three different input distances.

The Depolarization Block Test determines whether a model enters depolarization block in response to a prolonged and high-intensity somatic current stimulus and also determines the minimal current required to elicit this behavior. CA1 PCs respond to somatic current injections of increasing intensity with an increasing number of action potentials until a threshold is reached. For current intensities higher than the threshold, the cell does not fire over the whole period of the stimulus; instead, firing stops after some action potentials, and the membrane potential is sustained at some constant depolarized level for the rest of the stimulus. This phenomenon is known as depolarization block [20]. The test attempts to determine the threshold current by stimulating the soma of the model with 1000 ms long square current pulses of increasing amplitudes from 0 to 1.6 nA in 0.05 nA steps. The model is considered to exhibit depolarization block if the largest number of action potentials is not displayed at the highest current amplitude tested and if a higher current intensity exists where the model does not fire during the last 100 ms of its voltage response. Those models that do not enter depolarization block at all get an error score of 100.

The Oblique Integration Test evaluates the signal integration properties of radial oblique dendrites by providing an increasing number of synchronous or asynchronous clustered synaptic inputs that mimic glutamate uncaging experiments.

The test evaluates the voltage threshold for dendritic spike initiation defined as the expected somatic depolarization at which a step-like increase in peak dV/dt occurs; the proximal threshold defined the same way as above, but including only those results in the statistics where the proximal part of the examined dendrite was stimulated; distal threshold; degree of nonlinearity at threshold; suprathreshold degree of nonlinearity; peak derivative of somatic voltage at threshold; peak amplitude of somatic EPSP; time to peak of somatic EPSP; degree of nonlinearity in the case of asynchronous inputs. [17]

The test automatically selects a list of oblique dendrites that meet the criteria of being a terminal branch not further than 120 µm from the soma, then an increasing number of synaptic inputs are activated at the selected dendritic locations separately, while recording the local and somatic voltage response. To make the test mimic the glutamate uncaging experiments better, we put a single dendritic spine at the chosen dendritic location, which receives the synaptic input.

The synaptic weights for each selected dendritic location are automatically adjusted by the test using a binary search algorithm so that the threshold for dendritic spike generation is 5 synchronous inputs. Those dendritic locations where this first dendritic spike generates a somatic action potential, or where no dendritic spike can be evoked, are excluded from further analysis. We also modified the criteria for choosing dendritic location and weight -in the original test, the model should not fire any action potential for 5 inputs. We added that it should not fire at 6 inputs either, because somatic action potentials at 6 inputs interfere with the results and their interpretations. To reduce the probability of firing too early, we also gave a small (-0.1 nA) constant hyperpolarizing current injection to the soma.

In this test, we did not use the built-in NMDA receptor model of HippoUnit; instead, we tried several different NMDA receptor models found in the literature. To determine the right AMPA/NMDA ratio for each receptor model, we implemented the results of Otmakhov, Otmakhova, and Lisman [124]. We applied somatic voltage clamp at -65 mV, then recorded the voltage response of the cell when we gave no input to the cell, when we stimulated the cell with one single synaptic input on a proximal dendrite containing only AMPA current, and then when stimulated with a synaptic input with both AMPA and NMDA current in it. We subtracted the no-input trace from the other two cases as a correction, then took the integral of the corrected traces and calculated their ratio as follows:

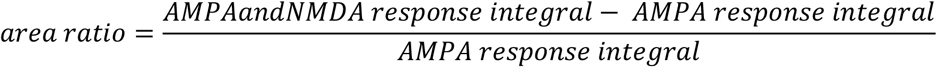

According to the article, this should result in a 3.7± 0.8 ratio in the Schaffer-collaterals. We used an AMPA/NMDA ratio for each NMDA receptor model that gives an area ratio within one standard deviation of the experimental result. To find the appropriate conductance ratio for a particular NMDA receptor model, we started from an AMPA/NMDA ratio of 0.5, and then if the area ratio was lower than 3.7± 0.8, we subtracted 0.1, and if it was higher, we added 0.1 until the area ratio got within 1 standard deviation of the experimental result. The model from Grunditz et al. could not be activated by one synaptic input sufficiently to get within the targeted range for the area ratio, and therefore, for this receptor model, we used the default 0.5 AMPA/NMDA ratio. Lastly, we created a new custom NMDA receptor model (see Methods above and Results) that fits our cell models best. The used AMPA/NMDA ratios with the corresponding area ratios can be seen in Table 5.

**Table 5:** The used AMPA/NMDA ratios for each NMDA receptor model with the corresponding area ratios.

| Receptor model | AMPA/NMDA ratio | Area Ratio |
| --- | --- | --- |
| Jahr-Stevens | 1 | 3.06 |
| Magee | 1.5 | 3.293 |
| Major | 0.4 | 3.475 |
| Gao | 0.4 | 3.116 |
| Otmakhova | 0.3 | 3.527 |
| Larkum | 0.25 | 3.436 |
| Ujfalussy | 0.5 | 3.874 |
| Tar (custom NMDA receptor) | 0.5 | 3.73 |

This version of HippoUnit, with the modifications made especially for this project and for validating models with dendritic spines, can be found at https://github.com/lucatar/hippounit/tree/hippounit_spine_project. Jupyter notebooks, target features, and the model can be found at https://github.com/lucatar/hippounit_CA1_demo.

## References

1. Almog M, Korngreen A. Is realistic neuronal modeling realistic? Journal of Neurophysiology. 2016; 116(5): 2180–2209.

2. Almog M, Korngreen A. A Quantitative Description of Dendritic Conductances and Its Application to Dendritic Excitation in Layer 5 Pyramidal Neurons. Journal of Neuroscience. 2014; 34(1): 182–196.

3. Hay E, Hill S, Schürmann F, Markram H, Segev I. Models of Neocortical Layer 5b Pyramidal Cells Capturing a Wide Range of Dendritic and Perisomatic Active Properties. PLOS Computational Biology. 2011; 7(7).

4. Schutter ED, Bower JM. An active membrane model of the cerebellar Purkinje cell. I. Simulation of current clamps in slice. Journal of Neurophysiology. 1994; 71(1): 375-400.

5. Poirazi P, Brannon T, Mel BW. Arithmetic of Subthreshold Synaptic Summation in a Model CA1 Pyramidal Cell. Neuron. 2003; 37(6): 977–987.

6. Migliore M, Hoffman DA, Magee JC, Johnston D. Role of an A-Type K+ Conductance in the Back-Propagation of Action Potentials in the Dendrites of Hippocampal Pyramidal Neurons. Journal of Computational Neuroscience. 1999; 7: 5–15.

7. Migliore R, Lupascu CA, Bologna LL, Romani A, Courcol JD, al. e. The physiological variability of channel density in hippocampal CA1 pyramidal cells and interneurons explored using a unified data-driven modeling workflow. PLOS Computational Biology. 2018; 14(9).

8. Káli S, Freund TF. Distinct properties of two major excitatory inputs to hippocampal pyramidal cells: a computational study. European Journal of Neuroscience. 2005; 22: 2027–2048.

9. Bezaire MJ, Raikov I, Burk K, Vyas D, Soltesz I. Interneuronal mechanisms of hippocampal theta oscillations in a full-scale model of the rodent CA1 circuit. eLife. 2016; 5: 1–106.

10. Markram H, Muller E, Ramaswamy S, M WR, Abdellah M, C AS, et al. Reconstruction and Simulation of Neocortical Microcircuitry. Cell. 2025; 163(2): 456–492.

11. Traub RD, Contreras D, Cunningham MO, Murray H, LeBeau FEN, Roopun A, et al. Single-Column Thalamocortical Network Model Exhibiting Gamma Oscillations, Sleep Spindles, and Epileptogenic Bursts. Journal of Neurophysiology. 2005; 93(4): 2194–2232.

12. Schneider CJ, Bezaire M, Soltesz I. Toward a full-scale computational model of the rat dentate gyrus. Frontiers in Neural Circuits. 2012; 6: 1–8.

13. Romani A, Antonietti A, Bella D, Budd J, Giacalone E, et a. Community-based reconstruction and simulation of a full-scale model of the rat hippocampus CA1 region. PLOS Biology. 2024; 22(11).

14. Mohácsi M, Török MP, Sáray S, Tar L, Farkas G, Káli S. Evaluation and comparison of methods for neuronal parameter optimization using the Neuroptimus software framework. PLOS Computational Biology. 2024; 20(12).

15. Geit WV, Gevaert M, Chindemi G, Rössert C, Courcol JD, Muller EB, et al. BluePyOpt: Leveraging Open Source Software and Cloud Infrastructure to Optimise Model Parameters in Neuroscience. Froniters in Neuroinformatics. 2016; 10.

16. Sáray S, Rössert CA, Appukuttan S, Migliore R, Vitale P, Lupascu CA, et al. HippoUnit: A software tool for the automated testing and systematic comparison of detailed models of hippocampal neurons based on electrophysiological data. PLOS Computational Biology. 2021; 17(1): 1–38.

17. Losonczy A, Magge JC. Integrative Properties of Radial Oblique Dendrites in Hippocampal CA1 Pyramidal Neurons. Neuron. 2006; 50(2): 291–307.

18. Magee JC, Cook EP. Somatic EPSP amplitude is independent of synapse location in hippocampal pyramidal neurons. Nature Neuroscience. 2000; 3: 895–903.

19. Golding NL, Kath WL, Spruston N. Dichotomy of Action-Potential Backpropagation in CA1 Pyramidal Neuron Dendrites. Journal of Neurophysiology. 2001; 86(6): 2998–3010.

20. Bianchi D, Marasco A, Limongiello A, Marchetti C, Marie H, Tirozzi B, et al. On the mechanisms underlying the depolarization block in the spiking dynamics of CA1 pyramidal neurons. Journal of Computational Neuroscience. 2012; 33: 207–225.

21. Zemankovics R, Káli S, Paulsen O, Freund TF, Hájos N. Differences in subthreshold resonance of hippocampal pyramidal cells and interneurons: the role of h-current and passive membrane characteristics. The Journal of Physiology. 2010; 588(12): 2109–2132.

22. Magee JC. Dendritic Hyperpolarization-Activated Currents Modify the Integrative Properties of Hippocampal CA1 Pyramidal Neurons. Journal of Neuroscience. 1998; 18(19): 7613–7624.

23. Lörincz A, Notomi T, Tamás G, Shigemoto R, Nusser Z. Polarized and compartment-dependent distribution of HCN1 in pyramidal cell dendrites. Nature Neuroscience. 2002; 5(11): 1185–93.

24. Lőrincz A, Nusser Z. Molecular Identity of Dendritic Voltage-Gated Sodium Channels. Science. 2010; 328(5980): 906–909.

25. Andrea L, Zoltán N. Molecular Identity of Dendritic Voltage-Gated Sodium Channels. Science. 2010; 328(5980): 906–909.

26. Coetzee WA, Amarillo Y, Chiu J, Chow A, Lau D, McCormack T, et al. Molecular Diversity of K+ Channels. Annals of the New York Academy of Sciences. 1999; 868(1): 233–255.

27. Misonou H, Mohapatra DP, Trimmer JS. Kv2.1: a voltage-gated k+ channel critical to dynamic control of neuronal excitability. Neurotoxicology. 2005; 26(5): 743–52.

28. Rudya B, McBain CJ. Kv3 channels: voltage-gated K+ channels designed for high-frequency repetitive firing. Trends in Neurosciences. 2001; 24(9): 517–526.

29. Metz AE, Spruston N, Martina M. Dendritic D-type potassium currents inhibit the spike afterdepolarization in rat hippocampal CA1 pyramidal neurons. The Journal of Physiology. 2007; 581(1): 175–187.

30. Storm JF. Potassium currents in Hippocampal pyramidal cells. Progress in Brain Research. 1990; 83.

31. Storm JF. Temporal integration by a slowly inactivating K+ current in hippocampal neurons. Nature. 1988; 336.

32. Hönigsperger C, Marosi M, Murphy R, Storm JF. Dorsoventral differences in Kv7/M-current and its impact on resonance, temporal summation and excitability in rat hippocampal pyramidal cells. Journal of Physiology. 2015; 593(7): 1551–1580.

33. Shah MM, Migliore M, Brown DA. Differential effects of Kv7 (M-) channels on synaptic integration in distinct subcellular compartments of rat hippocampal pyramidal neurons. Journal of Physiology. 2011; 589(24): 6029–6038.

34. Hoffman DA, Magee JC, Colbert CM, Johnston D. K+ channel regulation of signal propagation in dendrites of hippocampal pyramidal neurons. Nature. 1997; 387: 869–875.

35. Kerti K, Lorincz A, Nusser Z. Unique somato-dendritic distribution pattern of Kv4.2 channels on hippocampal CA1 pyramidal cells. European Journal of Neuroscience. 2011; 35(1): 66–75.

36. Andrew W. Varga LLY, Anderson AE, Schrader LA, Wu GY, Gatchel JR, Johnston D, et al. Calcium–Calmodulin-Dependent Kinase II Modulates Kv4.2 Channel Expression and Upregulates Neuronal A-Type Potassium Currents. Journal of Neuroscience. 2004; 24(14): 3643–3654.

37. Lindroos R, Dorst MC, D. K, Filipovic M, Keller D, Ketzef M, et al. Basal Ganglia Neuromodulation Over Multiple Temporal and Structural Scales—Simulations of Direct Pathway MSNs Investigate the Fast Onset of Dopaminergic Effects and Predict the Role of Kv4.2. Frontiers in Neural Circuits. 2018; 12.

38. Siwiec M, Kusek M, Sowa JE, Tokarski K, Hess G. 5-HT7 receptors increase the excitability of hippocampal CA1 pyramidal neurons by inhibiting the A-type potassium current. Neuropharmacology. 2020; 177.

39. Jerng HH, Pfaffinger PJ. Modulatory mechanisms and multiple functions of somatodendritic A-type K+ channel auxiliary subunits. Frontiers in Cellular Neuroscience. 2014; 8.

40. Kise Y, Kasuya G, Okamoto HH, Yamanouchi D, Kobayashi K, Kusakizako T, et al. Structural basis of gating modulation of Kv4 channel complexes. Nature. 2021; 599: pages158–164.

41. Murphy JG, Gutzmann JJ, Lin L, Hu J, Petralia RS, Wang YX, et al. R-type voltage-gated Ca2+ channels mediate A-type K+ current regulation of synaptic input in hippocampal dendrites. Cell Reports. 2022; 38(3).

42. McNaughton NCL, Randall AD. Electrophysiological properties of the human N-type Ca2+ channel: I. Channel gating in Ca2+, Ba2+ and Sr2+ containing solutions. Neuropharmacology. 1997; 36(7): 895–915.

43. Bowden SEH, Fletcher S, Loane DJ, Marrion NV. Somatic Colocalization of Rat SK1 and D class (Cav 1.2) L-type Calcium Channels in Rat CA1 Hippocampal Pyramidal Neurons. Journal of Neuroscience. 2001; 21.

44. Marrion NV, Tavalin SJ. Selective activation of Ca2+-activated K+ channels by co-localized Ca2+ channels in hippocampal neurons. Nature. 1998; 395: 900–905.

45. Magee J, Hoffman D, Colbert C, Johnston D. Electrical and Calcium Signaling in Dendrites of Hippocampal Pyramidal Neurons. Annual Review of Physiology. 1998; 60: 327–346.

46. Bell DC, Butcher AJ, Berrow NS, Page KM, Brust PF, Nesterova A, et al. Biophysical Properties, Pharmacology, and Modulation of Human, Neuronal L-Type (α1D, CaV1.3) Voltage-Dependent Calcium Currents. Journal of Physiology. 2001; 85(2): 816–827.

47. Randall AD, Tsien RW. Contrasting biophysical and pharmacological properties of T-type and R-type calcium channels. Neuropharmacology. 1997; 36(7): 879–893.

48. McKay BE, McRory JE, Molineux ML, Hamid J, Snutch TP, Zamponi GW, et al. Cav3 t-type calcium channel isoforms differentially distribute to somatic and dendritic compartments in rat central neurons. European Journal of Neuroscience. 2006; 24(9): 2581–2594.

49. Bloodgood BL, Sabatini BL. Nonlinear Regulation of Unitary Synaptic Signals by CaV2.3 Voltage-Sensitive Calcium Channels Located in Dendritic Spines. Neuron. 2007; 53(2): 249–260.

50. Yasuda R, Sabatini BL, Svoboda K. Plasticity of calcium channels in dendritic spines. Nature Neuroscience. 2003; 6: 948–955.

51. Foehring RC, Mermelstein PG, Song WJ, Ulrich S, Surmeier DJ. Unique Properties of R-Type Calcium Currents in Neocortical and Neostriatal Neurons. Journal of Neurophysiology. 2000; 84(5): 2225–2236.

52. Gu N, Vervaeke K, Storm JF. BK potassium channels facilitate high-frequency firing and cause early spike frequency adaptation in rat CA1 hippocampal pyramidal cells. Journal of Physiology. 2007; 580(3): 859–882.

53. Maurice N, Mercer J, Chan CS, Hernandez-Lopez S, Held J, Tkatch T, et al. D2 Dopamine Receptor-Mediated Modulation of Voltage-Dependent Na+ Channels Reduces Autonomous Activity in Striatal Cholinergic Interneurons. Journal of Neuroscience. 2004; 24(46): 10289–10301.

54. Ballesteros-Merino C, Lin M, Wu WW, Ferrandiz-Huertas C, Cabañero MJ, Watanabe M, et al. Developmental profile of SK2 channel expression and function in CA1 neurons. Hippocampus. 2012; 22(6): 1467–1480.

55. Hirschberg B, Maylie J, Adelman JP, Marrion NV. Gating of Recombinant Small-Conductance Ca-activated K+ Channels by Calcium. Journal of General Physiology. 1998; 111(4): 565–581.

56. King B, Rizwan AP, Asmara H, Heath NC, Engbers JDT, Dykstra S, et al. IKCa Channels Are a Critical Determinant of the Slow AHP in CA1 Pyramidal Neurons. Cell Reports. 2015; 11(2): 175 –182.

57. Traub RD, Wong RK, Miles R, Michelson H. A model of a CA3 hippocampal pyramidal neuron incorporating voltage-clamp data on intrinsic conductances. 635-650. 1991; 66(2): Journal of Physiology.

58. Bernard C, Johnston D. Distance-dependent modifiable threshold for action potential back-propagation in hippocampal dendrites. Journal of Neurophysiology. 2003; 90(3): 1807–16.

59. Vivo RSNECDCtNCI. Valerio Francioni; M. T. Harnett. Neuroscience. 2022; 489: 185–199.

60. Terbe D, Szoboszlay M, Z. Nusser SK. Reliable estimation of neuronal biophysical parameters from whole-cell recordings using model simulations and probabilistic inference. In Society for Neuroscience; 2022; San Diego.

61. Pierre F. Apostolides Admcgkcbjcm. atAxonal Filtering Allows Reliable Output during Dendritic Plateau-Driven Complex Spiking in CA1 Neurons. Neuron. 2016; 89(4): 770–783.

62. Spruston N. Pyramidal neurons: dendritic structure and synaptic integration. Nature Reviews Neuroscience. 2008; 9(3): 206–221.

63. Jahr CE, Stevens CF. Voltage dependence of NMDA-activated macroscopic conductances predicted by single-channel kinetics. Journal of Neuroscience. 1990; 10(9): 3178–3182.

64. Nonna A. Otmakhova NOaJEL. Pathway-Specific Properties of AMPA and NMDA-Mediated Transmission in CA1 Hippocampal Pyramidal Cells. Journal of Neuroscience. 2002; 22(4): 1199–1207.

65. Major G, Polsky A, Denk W, Schiller J, Tank DW. Spatiotemporally Graded NMDA Spike/Plateau Potentials in Basal Dendrites of Neocortical Pyramidal Neurons. Journal of Neurophysiology. 2008; 99(5): 2584–2601.

66. Ujfalussy BB, Makara JK. Impact of functional synapse clusters on neuronal response selectivity. Nature Communications. 2020; 11.

67. Grunditz Å, Holbro N, Tian L, Zuo Y, Oertner TG. Spine Neck Plasticity Controls Postsynaptic Calcium Signals through Electrical Compartmentalization. Journal of Neuroscience. 2008; 28(50): 13457–13466.

68. Larkum ME, Nevian T, Sandler M, Polsky A, Shiller J. Synaptic Integration in Tuft Dendrites of Layer 5 Pyramidal Neurons: A New Unifying Principle. Science. 2009; 325(5941): 756–760.

69. Andrásfalvy BK, Magee JC. Distance-Dependent Increase in AMPA Receptor Number in the Dendrites of Adult Hippocampal CA1 Pyramidal Neurons. Journal of Neuroscience. 2001; 21(23): 9151–9159.

70. Gao C, Gill MB, Tronson NC, Guedea AL, Guzmán YF, Huh KH, et al. Hippocampal NMDA receptor subunits differentially regulate fear memory formation and neuronal signal propagation. Hippocampus. 2010; 20(9): 1072–1082.

71. Guan D, Lee JCF, Higgs MH, Spain WJ, Foehring RC. Functional Roles of Kv1 Channels in Neocortical Pyramidal Neurons. Journal of Neurophysiology. 2007; 97(3): 1931–1940.

72. Liu PW, Bean BP. Kv2 Channel Regulation of Action Potential Repolarization and Firing Patterns in Superior Cervical Ganglion Neurons and Hippocampal CA1 Pyramidal Neurons. Journal of Neuroscience. 2014; 34(14): 4991–5002.

73. Metz AE, Spruston N, Martina M. Dendritic D-type potassium currents inhibit the spike afterdepolarization in rat hippocampal CA1 pyramidal neurons. Journal of Physiology. 2007;: 175-187.

74. Shah MM, Migliore M, Brown DA. Differential effects of Kv7 (M-) channels on synaptic integration in distinct subcellular compartments of rat hippocampal pyramidal neurons. Journal of Physilogy. 2011; 589(24): 6029–6038.

75. Cai X, Liang CW, Muralidharan S, Kao JPY, Tang CM, Thompson SM. Unique Roles of SK and Kv4.2 Potassium Channels in Dendritic Integration. Neuron. 2004; 44(2): 351–364.

76. Gasparini S, Losonczy A, Chen X, Johnston D, Magee JC. Associative pairing enhances action potential back-propagation in radial oblique branches of CA1 pyramidal neurons. Journal of Physiology. 2007; 580(3): 787–800.

77. Vogalis F, Storm JF, Lancaster B. SK channels and the varieties of slow after-hyperpolarizations in neurons. European Journal of Neuroscience. 2003; 18(12): 3155–3166.

78. Stocker M. Ca2+-activated K+ channels: molecular determinants and function of the SK family. Nature Reviews Neuroscience. 2004; 5: 758–770.

79. Tsay D, Yuste R. On the electrical function of dendritic spines. Trends in Neuroscience. 2004; 27(2): 77–83.

80. Yuste R. Dendritic spines and distributed circuits. Neuron. 2011; 71(5): 772–81.

81. Jan Tønnesen UVN. Dendritic Spines as Tunable Regulators of Synaptic Signals. Froniters in Psychiatry. 2016; 7(101).

82. Matsuzaki M, Ellis-Davies GCR, Nemoto T, Miyashita Y, Iino M, Kasai H. Dendritic spine geometry is critical for AMPA receptor expression in hippocampal CA1 pyramidal neurons. Nature Neuroscience. 2001; 4: 1086–1092.

83. Rapp M, Yarom Y, Segev I. The Impact of Parallel Fiber Background Activity on the Cable Properties of Cerebellar Purkinje Cells. Neural Computation. 1992; 4(4): 518–533.

84. Cornejo VH, Ofer N, Yuste R. Voltage compartmentalization in dendritic spines in vivo. Science. 2022; 375(6576): 82–86.

85. Yuste R. Electrical compartmentalization in dendritic spines. Annual Review of Neuroscience. 2013; 36: 429–449.

86. Lucy M. Palmer GJS. Membrane Potential Changes in Dendritic Spines during Action Potentials and Synaptic Input. Journal of Neuroscience. 2009; 29(21): 6897–6903.

87. Guy E, B. VM, Guilherme TS, Yair D, Ruth BP, Javier D, et al. Human Cortical Pyramidal Neurons: From Spines to Spikes via Models. Frontiers in Cellular Neuroscience. 2018; 12.

88. Zhang Y, He G, Ma L, Liu X, Hjorth JJJ, Kozlov A, et al. A GPU-based computational framework that bridges neuron simulation and artificial intelligence. Nature Communications. 2023; 14.

89. Lagache T, Jayant K, Yuste R. Electrodiffusion models of synaptic potentials in dendritic spines. Journal of Computational Neuroscience. 2019; 47(1): 77–89.

90. Druckmann S, Banitt Y, Gidon A, Schürmann F, Markram H, Segev I. A novel multiple objective optimization framework for constraining conductance-based neuron models by experimental data. Froniters in Neuroscience. 2007; 1(1): 7–18.

91. Hayes RD, Byrne JH, Cox SJ, Baxter DA. Estimation of single-neuron model parameters from spike train data. Neurocomputing. 2005; 65-66: 517–529.

92. Golding NL, Jung Hy, Mickus T, Spruston N. Dendritic Calcium Spike Initiation and Repolarization Are Controlled by Distinct Potassium Channel Subtypes in CA1 Pyramidal Neurons. Journal of Neuroscience. 1999; 19(20): 8789–8798.

93. Megiás M, Emri Z, Freund TF, Gulyás AI. Total number and distribution of inhibitory and excitatory synapses on hippocampal CA1 pyramidal cells. Neuroscience. 2001; 102(3): 527–540.

94. Hodgkin AL, Huxley AF. A quantitative description of membrane current and its application to conduction and excitation in nerve. Journal of Physiology. 1952; 117(4): 500–544.

95. Borg-Graham L. Modelling the non-linear conductances of excitable membranes. Cellular Neurobiology: A Practical Approach. 1991; 13: 247–275.

96. Mitrovic N, Jr. Alg, Horn R. Independent Versus Coupled Inactivation in Sodium Channels: Role of the Domain 2 S4 Segment. Journal of General Physiology. 1998; 111(3): 451–462.

97. Pan AC, Cuello LG, Perozo E, Roux B. Thermodynamic coupling between activation and inactivation gating in potassium channels revealed by free energy molecular dynamics simulations. Journal of General Physiology. 2011; 138(6): 571–580.

98. Cuello LG, Jogini V, Cortes DM, Pan AC, Gagnon DG, Dalmas O, et al. Structural basis for the coupling between activation and inactivation gates in K+ channels. Nature. 2010; 466: 272–275.

99. A L, A K. Markov modeling of ion channels: implications for understanding disease. Progress in Molecular Biology and Translational Science. 2014; 123: 1–21.

100. Siekmann I, Fackrell M, Crampin EJ, Taylor P. Modelling modal gating of ion channels with hierarchical Markov models. Proceedings of the Royal Society A. 2016; 472(2192).

101. Sansom MS, Ball FG, Kerry CJ, McGee R, Ramsey RL, Usherwood PN. Markov, fractal, diffusion, and related models of ion channel gating. A comparison with experimental data from two ion channels. Biophysical Journal. 1989; 56(6): 1229– 1243.

102. McKay BE, McRory JE, Molineux ML, Hamid J, Snutch TP, Zamponi GW, et al. CaV3 T-type calcium channel isoforms differentially distribute to somatic and dendritic compartments in rat central neurons. European Journal of Neuroscience. 2006; 24(9): 2581–2594.

103. Iftinca M, McKay BE, Snutch TP, McRory JE, Turner RW, Zamponi GW. Temperature dependence of T-type calcium channel gating. Neuroscience. 2006; 142(4): 1031–1042.

104. Chen X, Johnston D. Constitutively Active G-Protein-Gated Inwardly Rectifying K+ Channels in Dendrites of Hippocampal CA1 Pyramidal Neurons. Journal of Neuroscience. 2005; 25(15): 3787–3792.

105. Victoria NC, Velasco EMFd, Ostrovskaya O, Metzger S, Xia Z, Kotecki L, et al. G Protein-Gated K+ Channel Ablation in Forebrain Pyramidal Neurons Selectively Impairs Fear Learning. Biological Psychiatry. 2016; 80(10): 796–806.

106. Huang WC, Xiao S, Huang F, Harfe BD, Jan YN, Jan LY. Calcium-Activated Chloride Channels (CaCCs) Regulate Action Potential and Synaptic Response in Hippocampal Neurons. Neuron. 2012; 74(1): 179–192.

107. Gomez CD, Read J, Acharjee S, Pittman QJ. Early Life Inflammation Increases CA1 Pyramidal Neuron Excitability in a Sex and Age Dependent Manner through a Chloride Homeostasis Disruption. Journal of Neuroscinece. 2019; 39(37): 7244–7259.

108. Luca L. Bologna Rsdcmm. The EBRAINS NeuroFeatureExtract: An Online Resource for the Extraction of Neural Activity Features From Electrophysiological Data. Frontiers in Neuroinformatics. 2021; 15.

109. Harnett MT, Makara JK, Spruston N, Kath WL, Magee JC. Synaptic amplification by dendritic spines enhances input cooperativity. Nature. 2012; 491: 599–602.

110. Borg-Graham LJ. Interpretations of Data and Mechanisms for Hippocampal Pyramidal Cell Models. In Cerebral Cortex. New York: Plenum Press; 1999. p. 19–138.

111. Colbert CM, Magee JC, Hoffman DA, Johnston D. Slow Recovery from Inactivation of Na+ Channels Underlies the Activity-Dependent Attenuation of Dendritic Action Potentials in Hippocampal CA1 Pyramidal Neurons. Journal of Neuroscience. 1997; 17(17): 6512–6521.

112. Jung HY, Mickus T, Spruston N. Prolonged Sodium Channel Inactivation Contributes to Dendritic Action Potential Attenuation in Hippocampal Pyramidal Neurons. Journal of Neuroscience. 1997; 17(17): 6639–6646.

113. Martina M, Jonas P. Functional differences in Na+ channel gating between fast-spiking interneurones and principal neurones of rat hippocampus. The Journal of Physiology. 1997; 505(3): 593–603.

114. Chen X, Johnston D. Properties of single voltage-dependent K+ channels in dendrites of CA1 pyramidal neurones of rat hippocampus. Journal of Physiology. 2004;: 187–203.

115. Johnston D, Hoffman DA, Magee JC, Poolos NP, Watanabe S, Colbert CM, et al. Dendritic potassium channels in hippocampal pyramidal neurons. Journal of Physiology. 2000;: 75–81.

116. Kim J, Wei DS, Hoffman DA. Kv4 potassium channel subunits control action potential repolarization and frequency-dependent broadening in rat hippocampal CA1 pyramidal neurones. Journal of Physiology. 2005; 569(1): 41–57.

117. Kerti K, Lorincz A, Nusser Z. Unique somato-dendritic distribution pattern of Kv4.2 channels on hippocampal CA1 pyramidal cells. European Journal of Neuroscience. 2011; 35(1): 66–75.

118. Golomb D, Donner K, Shacham L, Shlosberg D, Amitai Y, Hansel D. Mechanisms of Firing Patterns in Fast-Spiking Cortical Interneurons. PLOS Computational Biology. 2007; 3(8).

119. Chen X, Johnston D. Properties of single voltage-dependent K+ channels in dendrites of CA1 pyramidal neurones of rat hippocampus. Journal of Physiology. 2004;: 187-203.

120. McDermott CM, Hardy MN, Bazan NG, Magee JC. Sleep deprivation-induced alterations in excitatory synaptic transmission in the CA1 region of the rat hippocampus. Journal of Physiology. 2006; 570(3): 553–565.

121. Carnevale NT, Hines ML. The Neuron Book New York, NY, USA: Cambridge University Press; 2009.

122. Friedrich P, Vella M, Gulyás AI, Freund TF, Káli S. A flexible, interactive software tool for fitting the parameters of neuronal models. Frontiers in Neuroinformatics. 2014; 8(63).

123. Sivagnanam S, Majumdar A, Yoshimoto K, Astakhov V, Bandrowski A, Martone ME, et al. Introducing the Neuroscience Gateway. CEUR Workshop Proceedings. 2013; 993.

124. Otmakhova NA, Otmakhov N, Lisman JE. Pathway-Specific Properties of AMPA and NMDA-Mediated Transmission in CA1 Hippocampal Pyramidal Cells. The Journal of Neuroscience. 2002; 22(4): 1199–1207.

125. Migliore M. Modeling the attenuation and failure of action potentials in the dendrites of hippocampal neurons. Biophysical Journal. 1996; 71(5): 2394–2403.

126. Zühlke RD, Pitt GS, Deisseroth K, Tsien RW, Reuter H. Calmodulin supports both inactivation and facilitation of L-type calcium channels. Nature. 1999; 399: 159–162.

127. Segev I, Rall W. Computational study of an excitable dendritic spine. Journal of Neurophysiology. 1988; 60(2): 499–523.

